# Prolonged development of tonotopic tuning in human auditory cortex

**DOI:** 10.64898/2026.04.20.719686

**Authors:** Oyindamola Ogunlade, Jesse Gomez

## Abstract

Audition is a fundamental sense, underlying critical human behaviors such as communication and recognition. Despite its importance, how the tuning and organization of receptive fields mature from childhood to adulthood in auditory cortex has not been directly measured. Through a gamified neuroimaging approach using functional MRI, we model population receptive field (pRF) tuning for frequency across human auditory cortex in both children and adults. In the same participants, we behaviorally quantify detection thresholds for different frequencies embedded in noise to understand how the functional development of human auditory cortex drives behavior. We find that while the tonotopic organization of pRFs is qualitatively present in early childhood, there is a protracted increase in the representation of low frequencies in tonotopic maps of auditory cortex. This maturation of pRF tuning appears to drive basic auditory behaviors, relating to tone detection thresholds across participants. We also observe protracted development in secondary auditory regions, offering evidence for an anatomically-predictable tonotopic map posterior to Heschl’s Gyrus. These data provide a new avenue for studying the development of audition in the human brain and lay important groundwork for understanding atypical development in auditory processing disorders.

## Introduction

The ability to effectively navigate complex auditory environments plays an integral role in daily life, yet how low-level^1–5^ and complex^6–11^ auditory features are encoded across primary and non-primary auditory cortex remains unclear, let alone how auditory representations develop during childhood. Comparatively, it has been well-established that the retinotopic visual field maps of visual space^12–15^ are organized across cortex such that they follow the underlying macroanatomy, with the boundaries between visual field maps often occurring along cortical folds across the visual processing hierarchy^16–18^. Developmental work has uncovered that visual cortex shows protracted maturation in how it pools information from visual space^19^, developing hierarchically with later-stage regions showing the most prolonged development. In contrast, the field of auditory neuroscience lacks consensus regarding the arrangement of tonotopic maps in superior temporal cortex, and how their tuning develops during critical childhood years. One possibility is that by studying auditory cortex development, we can identify which regions show developmental dynamics versus stability, potentially gaining insight into their hierarchical arrangement in human cortex.

Much of what is understood about the functional organization of auditory cortex results from micro-electrode work in animals^20–23^ and early cytoarchitectonic studies in humans^24–26^. Such work has established the tonotopic organization of the mammalian auditory cortex in which neurons are arranged according to a frequency gradient that is inherited from the cochlea. In macaques, auditory cortex consists of a heavily myelinated primary “core” which processes and relays low-level auditory information to secondary “belt” and “parabelt” regions via parallel and serial connections^27^. The core contains tonotopic maps whose frequency reversals delineate subfields A1, R and RT. Most notably, the low frequency preferring region of a high-low-high frequency gradient marks the boundary between A1 and R. Neuroimaging work has long confirmed that human primary auditory cortex similarly comprises a mirror-symmetric high-low-high frequency gradient^4,28,29^. However, unlike the monkey auditory cortex whose belt and parabelt have been further parcellated into functionally defined auditory fields^27,20,30^ a consensus on the tonotopic organization of human auditory cortex beyond the core has yet to be reached^5,29,31,32^. This is due in part to disagreement in the field over the orientation of the core in relation to HG. A “classical” interpretation positions the core parallel to the long axis of HG, and an “orthogonal” framework places the core perpendicular to HG and extending anteriorly and posteriorly into the planum polare (PP) and planum temporale (PT) respectively^29^.

It is unclear how the computations of neurons within human auditory cortex change across development. Behavioral work suggests that the processing of simple auditory features continues to develop into adulthood. Auditory sensitivity to pure tones has been shown to change across development in a frequency-specific manner^33,34^. While sensitivity to high attfrequencies reaches maturity by 5 years old, thresholds for perceiving low frequencies may continue to develop into late childhood^33^, with tone-in-noise detection showing a similarly protracted development^35^. It has been proposed that this improvement is related to changes in the “filter” properties of the auditory system^35,36^. This behavioral evidence for the protracted development of auditory processing is underscored by structural changes occurring in the auditory system throughout childhood. For example, an increase in mastoid air-cell volume from childhood to adulthood^37^ may promote the transduction of low frequency sound waves^35,37^. Further, late childhood (5-12 years old) is characterized by axonal maturation which supports more efficient relay between different subregions of auditory cortex, fine-tuning the processing of auditory stimuli^38^. These findings thus suggest that there should be protracted development of the neural tuning properties of human auditory cortex across childhood. While there is evidence for the maturation of tonotopic organization of auditory evoked potential components^39^, hurdles have prevented the mapping of auditory receptive fields in childhood populations given the difficulty of neuroimaging younger participants. This has left unclear how functional development unfolds across the auditory processing hierarchy. Establishing this developmental model has important implications for our understanding of neurodevelopmental disorders involving deficits in auditory processing, including autism^41–44^ and auditory processing disorder^44–46^.

Developmental work in the visual domain^18,19,47^ and across the cortical sheet more broadly^48^ has demonstrated that sensorimotor systems develop hierarchically. Primary sensorimotor regions like V1 or M1 are more heavily myelinated than downstream regions^49,50^, and primary regions reach maturity sooner in development than secondary and higher-level regions of the processing hierarchy^51–53,19^. In this developmental framework, we would expect primary auditory regions like the auditory core (i.e., tonotopic maps hA1, hR, and hRT) to be earlier developed than higher-order auditory regions like the belt or parabelt. However, when one considers the competing organizational frameworks of human auditory cortex, we can see that they make distinct predictions regarding where functional development should be most dramatic. The orthogonal model conceives of the two tonotopic gradients straddling HG as primary auditory cortex^4,54^, given that these two tonotopic maps share a low frequency representation like the macaque model. However, structural and cytoarchitectonic mapping suggest that only the most medial portion of HG is heavily myelinated^55^, and cytoarchitectonic zones^56^ do not map neatly onto the anterior and posterior functional split of the orthogonal model. This has led some to propose a “classical” organization framework, which proposes the auditory core should be mapped in humans parallel with the long axis of HG, along its anterior side^29^. This model thus places secondary maps like the lateral belt on the posterior aspect of HG, where the orthogonal model places hA1. Given this mutually exclusive assignment of posterior HG as either belonging to primary (hA1) or secondary (belt) auditory cortex, determining how functional development of posterior and anterior aspects of HG compare to one another can help us adjudicate between organizational frameworks. Mapping the development of auditory cortex thus provides us a foothold for better understanding its hierarchical organization.

To break ground in pediatric neuroimaging of human auditory cortex, we produced two gamified auditory experiments that engage younger children to both quantify auditory behavior and receptive field tuning in auditory cortex. The first game–*Noisy Nest*–measures detection thresholds of various frequencies with a tone-in-noise approach. This gamified task provides a behavioral metric that can be compared to neural responses modeled through functional magnetic resonance imaging (fMRI). To measure and model frequency tuning in auditory cortex, we designed a second game–*RoboTones*–a gamified population receptive field (pRF) mapping experiment optimized for pediatric fMRI adapted from prior approaches in adults^57^. The pRF mapping method is a stimulus encoding model that allows us to elucidate how auditory neural populations sample frequency or “cochleotopic” space^57,58^ with parameters that can be directly compared between children and adults. Because participants listened to similar pure-tone stimuli during fMRI which presents a concurrent source of ambient noise, the *Noisy Nest* tone-in-noise detection task’s inclusion of tones embedded in noise facilitates a more direct comparison between behavioral and neural responses.

Both children and adults completed behavioral testing and neuroimaging experiments described above. With these data, we: 1) determined whether there are qualitative differences across age groups in the tonotopic organization of auditory cortex, 2) quantified developmental differences in auditory pRF properties, and 3) modeled how pRF development of auditory cortex relates to differences between children and adults in auditory behavior. In our quantification of auditory pRF development in superior temporal cortex, we compared developmental effects in regions of cortex defined using both the classical and orthogonal frameworks. Lastly, we asked if functional development extends beyond Heschl’s Gyrus in higher-level auditory cortex of the planum temporale. While there is no current agreement on the precise topography of tonotopic maps in this part of cortex, we investigated whether shared cortical folding features beyond HG could be used to consistently identify additional tonotopic maps in order to measure their development.

## Results

### Tone-in-noise detection performance improves with age for low frequencies

We designed a gamified 3-alternative forced choice (3-AFC) task (*Noisy Nest*), adapted from a previous psychoacoustic study^35^ to assess tone-in-noise detection abilities in children (n = 32, 5-11 years old) and adults (n = 35, 21-34 years old). Tone-in-noise detection has not yet been assessed in children at low frequencies in the range of the human voice. Given the prolonged development of voice recognition abilities^59^ and prior pure-tone sensitivity mapping in children suggesting children are less sensitive to lower frequencies than adults^33^, we included a 150 Hz condition to test the hypothesis that children will require higher sound levels to detect a low frequency tone embedded in noise. We included the condition of noise to better match this behavioral task (completed outside of the scanner) with the environment of fMRI scanning which included ambient noise during stimulus mapping. Participants were prompted to select one of three auditory stimuli which contained a pure-tone masked by white noise (**Fig. 1a**). The white noise could either cover a broad frequency spectrum (flat condition) or be notched at the frequency of the embedded tone (notched condition) (**Fig. 1b and 1c**). The flat noise condition provides a reference to observe how tone detection is impacted when a notch is provided in children versus adults. The sound level of the tone was adjusted for each trial using an adaptive staircasing procedure^60^ (**Fig. 1d**) to derive for each participant their detection threshold (in decibels) for three distinct frequencies: 150 Hz, 2000 Hz, and 4000 Hz. We asked whether adults and children differed in their thresholds for detecting the tones across the two white noise conditions. Specifically, based on early psychoacoustic work^35,36^, we have the strong *a priori* hypothesis that children should show worse behavioral performance at the lowest frequency in the notched noise condition given that their perceptual auditory filters^61^ are more sensitive to neighboring noise.

**Figure 1:**
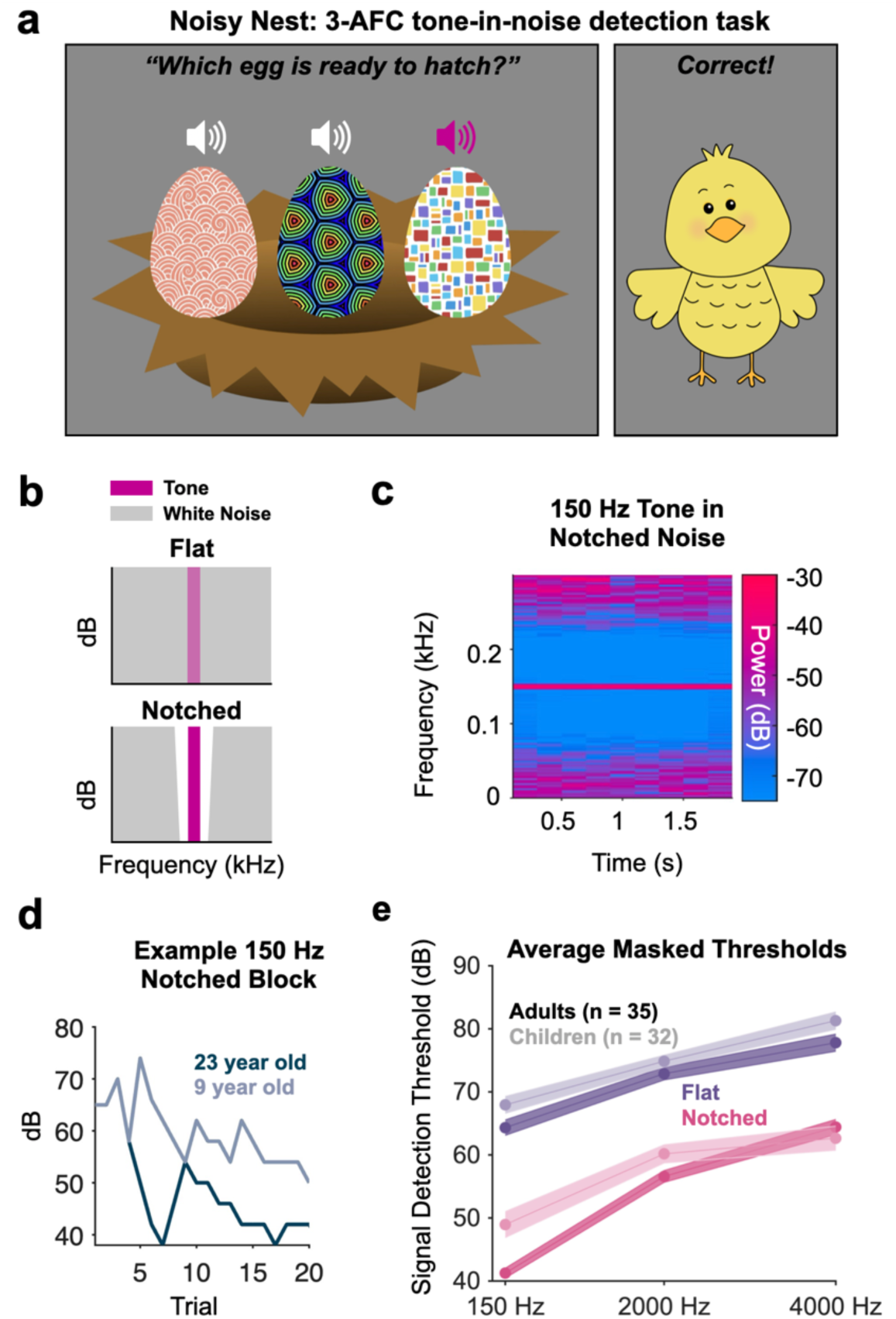
3-AFC tone-in-noise detection task and performance. **(a)** Left: Example trial of the Noisy Nest task. White volume icons indicate the eggs which emit white noise and the pink volume icon indicates the egg which emits the tone (150 Hz, 2000 Hz or 4000 Hz) embedded in white noise. Right: Correct feedback screen. **(b)** Visualization of the two white noise conditions: flat spectrum white noise (top) and white noise notched at the frequency of the tone (bottom). **(c)** Spectrogram of 150 Hz tone embedded in white noise notched at the same frequency. **(d)** Example block for 150 Hz tone embedded in notched spectrum noise in an example adult and child participant. **(e)** Signal detection thresholds averaged within adults (n = 35) and children (n = 32) for all conditions. Error bars represent standard deviation.

Both age groups demonstrated an increase in tone-in-noise detection thresholds with signal frequency in both white noise conditions (**Fig. 1e**). To investigate age group differences, a linear mixed-effects (LME) model was fit to detection thresholds with age group, frequency and white noise condition as fixed effects and a random intercept for participant. A significant main effect of frequency (F(2, 368) = 32.67, p < 0.001) and white noise (F(1, 368) = 187.14, p < 0.001) was observed, but not of age (F(1, 368) = 3.83, p = 0.051). Further, a significant interaction was found between frequency and white noise condition (F(2, 368) = 8.72, p < 0.001) as well as age, frequency and white noise condition (F(2, 368) = 3.67, p = 0.03). The significant three-way interaction was assessed with follow-up pairwise contrasts from the LME model assessing age group differences for each combination of frequency and white noise condition (p-values Bonferroni-corrected for multiple comparisons). These revealed lower detection thresholds in adults compared to children specifically for the 150 Hz notched white noise condition (estimated difference = 7.73, 95% CI [4.07, 11.38], F(1, 368) = 4.16, p < 0.001). These results suggest that children, like adults, can take advantage of less noise near the central frequency to aid in tone detection. However, adults benefit to a greater extent specifically at 150 Hz, suggesting children are more sensitive to nearby auditory noise at low frequencies.

### Tonotopic maps are observable in childhood auditory cortex and are qualitatively organized like adults

Adults (n = 25, 21-35 years old) and children (n = 25, 5-11 years old) completed the same tonotopic mapping procedure (*RoboTones*) in which they were presented with cartoon robots that moved across the screen, each emitting a pure frequency tone (**Fig. 2a**). Each tone was presented in a Morse-code pattern in random ascending and descending semi-sweeps^57^ (**Fig. 2b**). Participants responded with a button press when they perceived the oddball (tone repeat). Though head motion was significantly reduced in adults (mean frame-wise displacement 0.24 ± 0.12 mm) compared to children (0.47 ± 0.14 mm) (two-sample t-test: t(43) = −5.81, p < 0.001), no participants’ head motion exceeded our voxel size of 2 mm, and mean head motion was below the threshold of 0.5 mm mean framewise displacement used in prior work^62^. Importantly, as mentioned below, pRF model fits were similar in quality between children and adults. Furthermore, all children and adults completed 3 runs of tonotopic mapping with identical stimulus presentation across runs to enable averaging across runs and improve signal-to-noise ratio. We fit a 1-D Gaussian pRF at each vertex based on an established fitting procedure^57^ deriving a preferred frequency (μ) and bandwidth tuning estimate (σ). The pRF model performance was assessed by averaging the variance explained by the pRF model in a broad auditory cortex ROI including both the core and surrounding auditory cortex comprising the superior temporal plane and the superior temporal gyrus (**Fig. 2d, inset**). Variance explained by the pRF model was matched across age groups by excluding two children with the lowest pRF model fits and four adults with the highest pRF model fits in this ROI. As a result, there were no significant differences in variance explained by the pRF model in auditory cortex (**Fig. 2d**) between age groups (two-sample t-test: t(38) = 1.88, p = 0.07). This ensured that any developmental results observed did not derive from quality of model fit differences. The pRF model fits in an example adult and child (**Fig. 2c**) demonstrate high data quality across ages.

**Figure 2:**
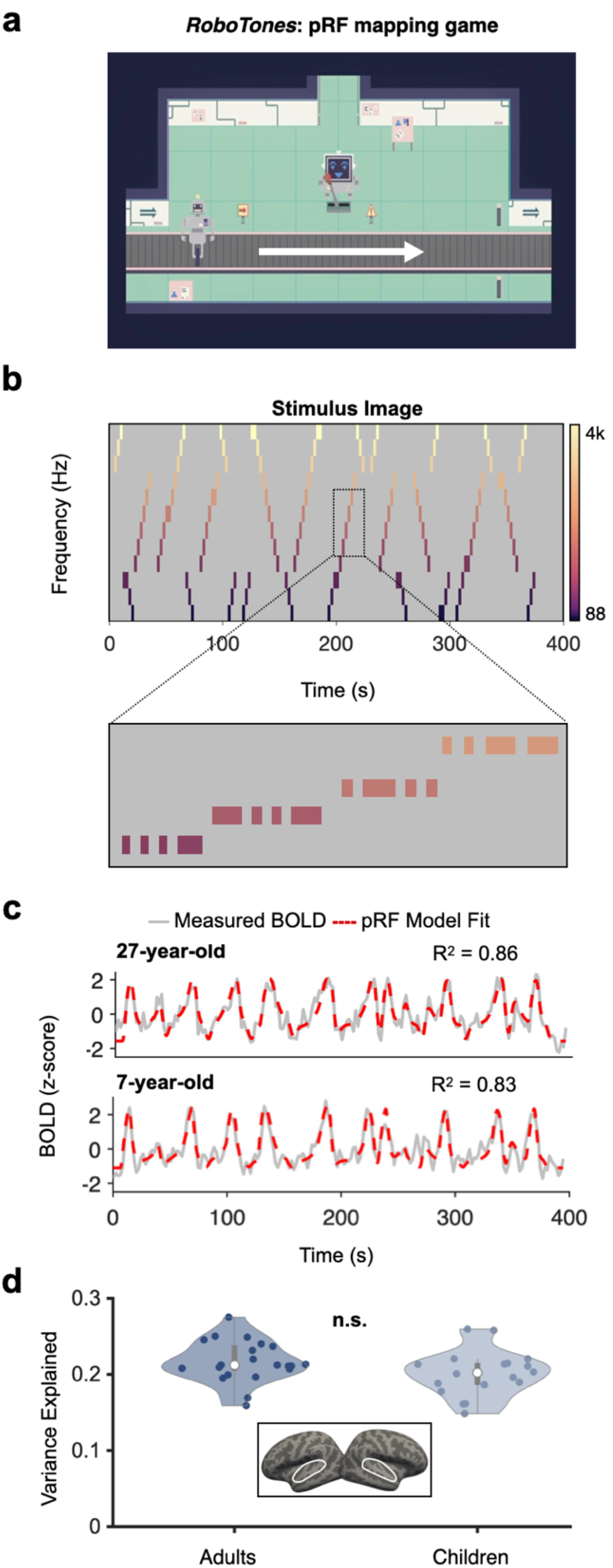
Tonotopic pRF mapping. **(a)** Adults (n = 25) and children (n = 25) underwent fMRI while playing RoboTones–a gamified tonotopic mapping task. Participants were presented with cartoon robots which moved across the screen on a conveyor belt, each emitting a pure frequency tone. Participants controlled a lever robot to teleport broken robots for repair. **(b)** Stimulus image of tonotopic mapping task indicating when each pure tone was presented throughout the run. Pure frequency tones were presented in random ascending and descending semi-sweeps from 88 Hz to 4000 Hz in half-octave steps. **(c)** The pRF predicted time course compared to the actual time course of a “HG” voxel in an example adult and child. **(d)** Variance explained within an auditory ROI comprising primary auditory cortex, the superior temporal plane and the superior temporal gyrus was matched between adults and children (two-sample t-test: t(37) = 1.68, p = 0.10).

Adults and children showed qualitatively similar tonotopic organization in their preferred frequency maps (**Fig. 3a**). When producing an average surface map of preferred frequency using cortex-based alignment, both children and adults demonstrate the expected low frequency representation centered on Heschl’s Gyrus (HG) with two diverging high frequency gradients extending posterior and anterior to the center of HG. As established in previous work^63^, HG was morphologically variable across individuals (**Fig. 3b)**, presenting as a single gyrus (left hemisphere: 20% of adults, 47% of children; right hemisphere: 8% of adults, 32% of children) or a duplicated gyrus (left hemisphere: 80% of adults, 53% of children; right hemisphere: 92% of adults, 68% of children). Despite this structural variability, HG is an anatomical landmark for the low frequency portion of the main mirror-symmetric map visible within individual participants (**Fig. 3b**), as this region was consistently centered on the peak of HG (in the case of a single HG) or its intermediate sulcus (in the case of a duplicated HG) across participants.

**Figure 3:**
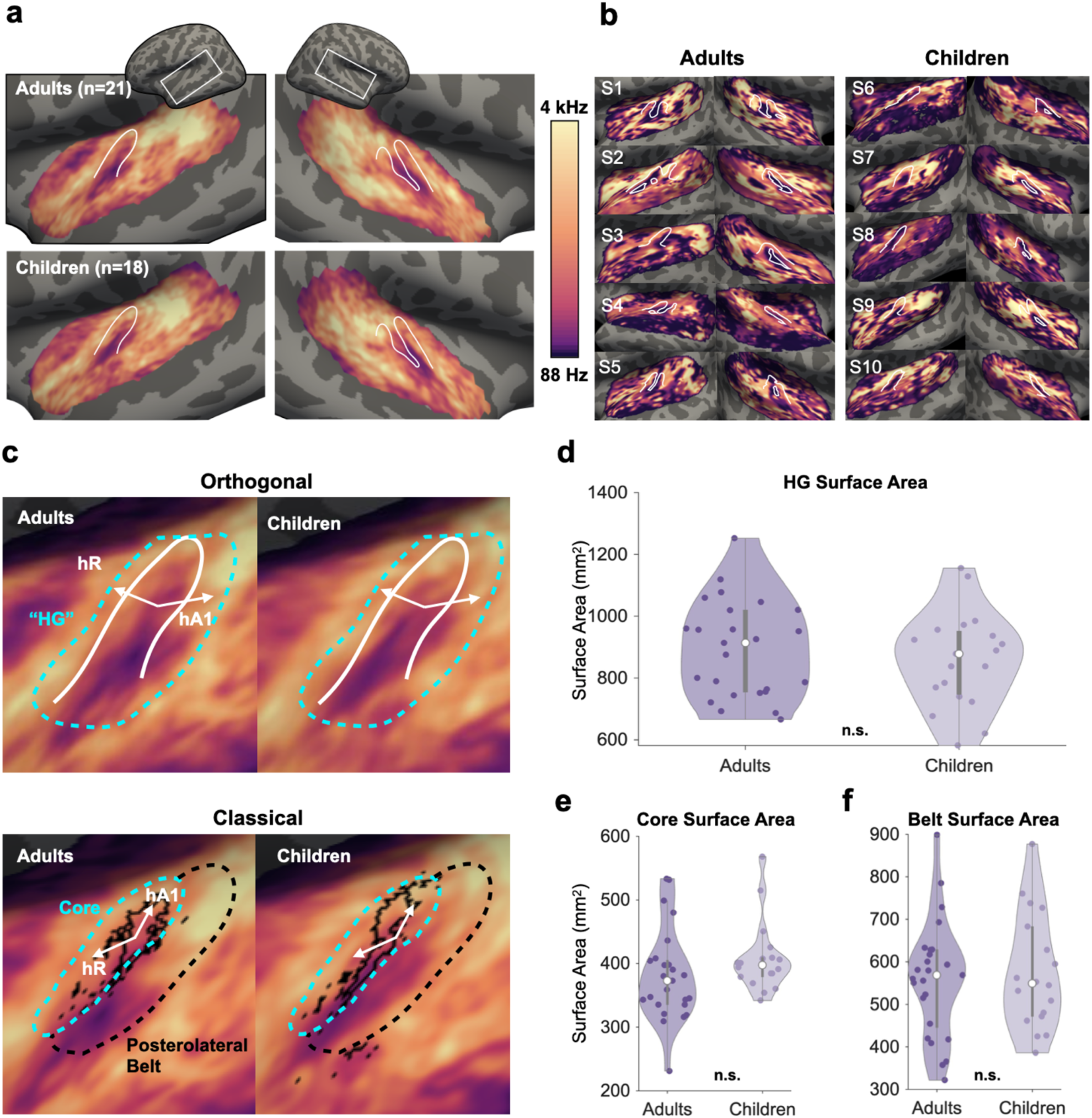
Tonotopic organization of the auditory core and posterolateral belt is qualitatively present in children and adults. **(a)** Preferred frequency (μ) maps in left and right auditory cortex averaged across n = 21 adults (top) and n = 18 children (bottom) and visualized on the fsaverage surface. Heschl’s Gyrus is outlined in white. **(b)** Preferred frequency maps projected onto the individual cortical surfaces of 5 example adults (left) and 5 example children (right) with Heschl’s Gyrus outlined in white. **(c)** Averaged preferred frequency maps in auditory cortex of adults (left) and children (right), wherein both groups demonstrate a mirror-symmetric high-low-high frequency gradient. Example definitions of the auditory core outlined in cyan according to “orthogonal” (top) and “classical” (bottom) orientations. Tonotopic gradients of subfields hA1 and hR are indicated by white arrows and label of top 10% myelin contrast vertices indicated by solid black line. The posterolateral belt has also been defined in the classical model (dotted black line) with its associated tonotopic gradient indicated by black arrow. **(d-f)** Violin plots depicting surface area in adults and children of the orthogonal “HG” ROI, the classical core ROI, and the belt ROI.

From this topography of frequency preference, we delineated on each participant’s cortical surface functional subdivisions (i.e., tonotopic maps) according to the two competing frameworks of auditory cortex organization. For the orthogonal model, we defined the primary auditory core as the pair of diverging, mirror-symmetric tonotopic gradients centered on HG (**Fig. 3c**, top). To avoid confusion with the parallel model, we refer to this orthogonal definition of the core as the “HG” ROI given that it spans the posterior and anterior halves of HG. Notably, this broad HG definition may contain some aspects of surrounding belt regions like CM/CL on the medial side^29^. For the classical model, the same expanse of cortex is divided differently. The core is instead defined on the anterior portion of HG encompassing a region of high intracortical myelination (**Fig. 3c**, bottom; Methods). We did not subdivide the core into its constituent fields (hA1, hR, hRT), keeping it instead as a conglomerate core ROI. To identify this cortical signature of myelination, we produced in each participant a map of the ratio of T1-weighted to T2-weighted images, an MR-based contrast thought to reflect the cortical density of myelin or neurite structures^64,65,50^. This contrast map was projected onto each participant’s cortical surface and thresholded to identify the top 10% highest contrast vertices near HG. The medial extent of the classical core was determined by the myelin-dense zone as done in previous work^29^, and its posterior/anterior extents were defined by the low-to-high frequency progression on the anterior HG. While the classical model defines the belt as a region surrounding the core in both anteromedial and posterolateral directions, we only defined the posterolateral belt here as it could be easily defined as the posterior frequency gradient on the opposing side of HG from the core (**Fig. 3c**, black arrow). While the classical and orthogonal ROIs delineated here occupy a similar cortical expanse (anterior and posterior halves of HG), they differ in where the primary auditory core is placed. The classical model places the core on the anterior half of HG, while the orthogonal model extends the core, specifically A1, onto the posterior half of HG.

We found no significant difference in mean surface area of any ROI. For the orthogonal HG ROI (**Fig. 3d**), there is no difference in surface area measured as either raw surface area (two-sample t-test: t(42) = 0.70, p = 0.49) or normalized by total cortical surface area (two-sample t-test: t(42) = 1.26, p = 0.21). For the classical model ROIs, the core between adults and children was also of similar surface area (two-sample t-test: t(41) = −1.34, p = 0.19; **Fig. 3e**). This effect does not change when normalizing surface area by total cortical surface area (adults mean percent whole brain surface area = 0.40%, children mean percent whole brain surface area = 0.41%, two-sample t-test: t(41) = −0.77, p = 0.45). Further, no significant difference in surface area was found in the posterolateral belt (raw surface area: t(41) = −0.38, p = 0.70; normalized surface area: t(41) = −0.08, p = 0.94; **Fig. 3f**).

### Auditory pRFs across HG retune to cover low frequency space with development

If adults show evidence of higher sensitivity to low frequency tones compared to children (**Fig. 1e**), do pRF properties reflect this developmental change? We compared pRF parameters—preferred frequency (μ) and bandwidth tuning (σ)—across age groups within the HG ROI, as well as the classically-defined core and belt ROIs, to quantify developmental differences in how auditory neural populations represent frequency. We began with the broader HG ROI which encompasses the two diverging tonotopic gradients centered on HG. In each participant, we derived the mean preferred frequency (μ) across all above-threshold vertices. Within the HG ROI, adults demonstrated significantly lower preferred frequency compared to children (two-sample t-test: t(37) = −2.61, p = 0.01; Fig. **4a**). To ensure this was not driven by differences in pRF model performance as a function of frequency across age groups, we performed an ANOVA and found no significant interaction between μ and variance explained (F(1, 35) = 3, p = 0.09) and no evidence that the relationship between μ and variance explained depends on age (F(1, 35) = 0.28, p = 0.6). We next examined the development of average bandwidth tuning (σ), age groups did not show any significant difference (two-sample t-test: t(37) = −0.2, p = 0.84; **Fig. 4b**). To get a finer grained understanding of pRF development, we asked whether bandwidth tuning may develop as a function of preferred frequency. We analyzed bandwidth tuning in a manner consistent with the behavioral data (**Fig. 1**) by defining a low frequency preference bin as μ < 150 Hz and a high frequency preference bin as μ > 2000 Hz and assigned voxels accordingly. We observed no significant difference in bandwidth tuning within low vs high frequency preferring voxels between age groups (two-way repeated-measures ANOVA, age x frequency preference interaction: F(1, 149) = 2.92, p = 0.09).

**Figure 4:**
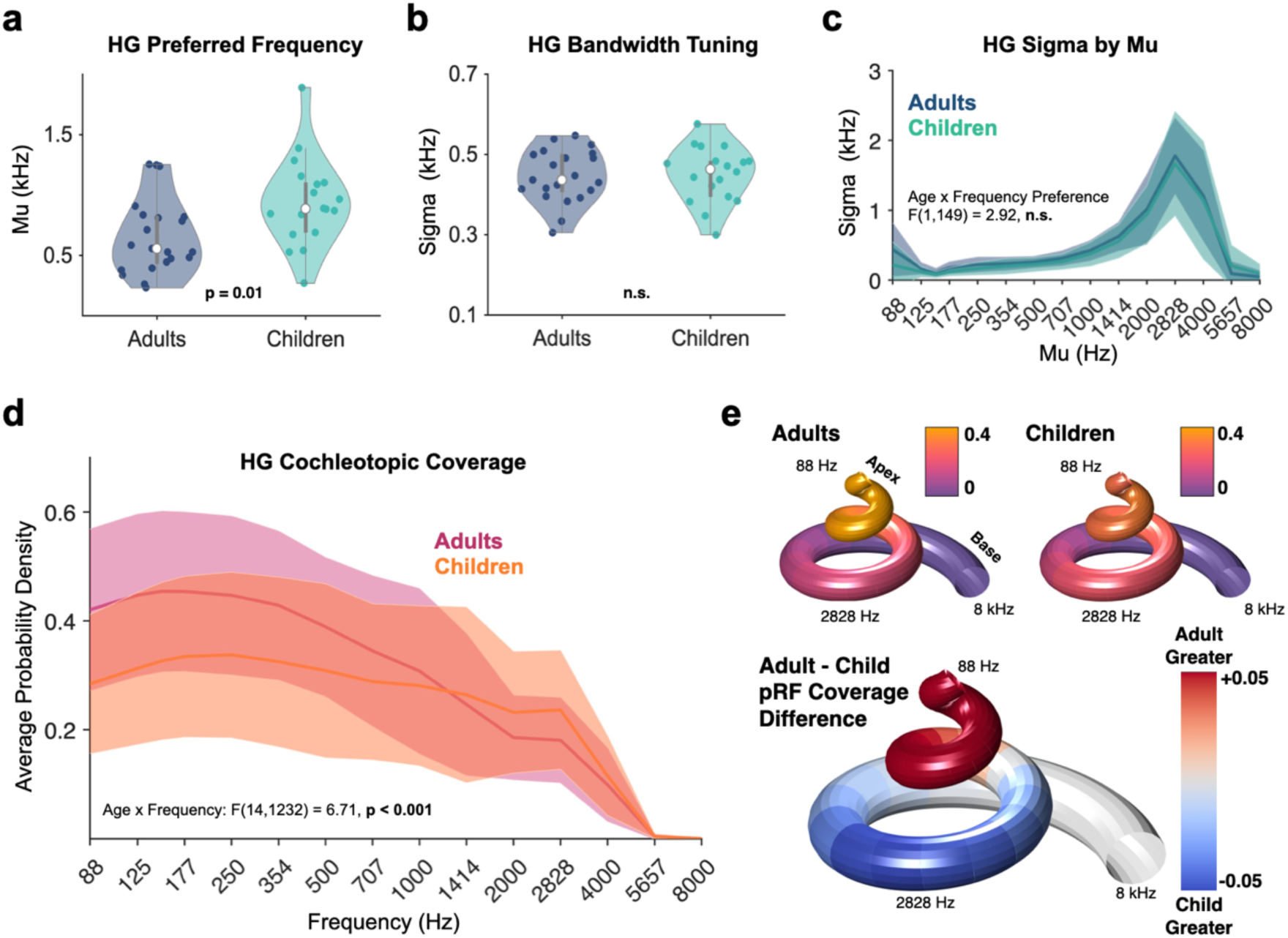
HG pRFs retune to more heavily cover low frequency space with development. **(a)** The average preferred frequency (μ) across pRF within the orthogonal HG ROI was determined for each participant. Group means denoted with white circles and standard deviation denoted with gray vertical lines. Adults showed significantly lower average μ in the HG ROI compared to children (t(37) = −2.61, p = 0.01). **(b)** The average pRF bandwidth tuning (σ) in each participant showed no developmental differences between groups (t(37) = −0.2, p = 0.84). **(c)** Histogram depicting mean pRF bandwidth as a function of frequency bin, separately plotted for children and adults, with shaded error bars denoting standard deviation. Within each participant’s HG ROI, pRFs were assigned to low μ (μ < 150 Hz) and high μ (μ > 2000 Hz) bins. The bandwidth tuning (σ) was averaged within these low μ and high μ voxels per participant. No difference in σ was observed across low μ and high μ voxels between adults and children (two-way repeated-measures ANOVA: age x frequency interaction: F(1, 149) = 2.92, p = 0.09). **(d)** The cochleotopic coverage of voxels within the HG ROI was determined for each participant and averaged within age groups. Adults and children demonstrated significantly different cochleotopic coverage patterns: ANOVA interaction of age and frequency, F(14, 1232) = 6.71, p < 0.001. **(e)** Visualization of cochleotopic coverage in adults (top left) and children (top right) on a model of the human cochlea. The difference of pRF coverage between adults and children (bottom) shows that adults demonstrate significantly more cochleotopic coverage than children at low frequencies while children have more coverage at higher frequencies.

Averaging pRF parameters within the HG ROI, while informative, is a reductive metric that does not capture the way pRFs sample cochleotopic input. To determine how densely auditory pRFs sample frequency space and how this may change across development, we compared cochleotopic coverage in this ROI between adults and children (**Fig. 4d**). Cochleotopic coverage was determined by reproducing each 1-D Gaussian within a participant in frequency space and then deriving the average pRF coverage value at each frequency. Cochleotopic coverage was normalized within each participant so that the resulting values range from 0 to a maximum of 1, allowing for direct comparison of cochleotopic coverage across participants with different numbers of vertices. The advantage of this approach is that it produces a more comprehensive and holistic model of population-level tuning within a cortical expanse for a given frequency. For visualization, all cochleotopic coverage curves were averaged across participants within an age group. A two-way repeated-measures ANOVA with frequency and age as the within- and between-subject factors respectively, revealed a main effect of age (F(1, 88) = 12.56, p < 0.001), a main effect of frequency (F(14, 1232) = 103.74, p < 0.001) and an interaction between age and frequency (F(14, 1232) = 6.71, p < 0.001). We visualized cochleotopic coverage differences on a simulated cochlea plot (**Fig. 4e**) where one can see that the adult HG ROI has a lower-frequency biased sampling of frequency space (heavier coverage of frequencies transduced near the apex of the cochlea) compared to children.

While these results suggested a protracted development of pRF tuning across Heschl’s Gyrus, we investigated to what extent the anteromedial and posterolateral halves of the diverging tonotopic gradients might develop distinctly. The orthogonal model places the hR map within the anteromedial portion and hA1 map in the anterolateral portion. The classical model, however, places the primary auditory core (the union of maps hA1, hR, and RT) on the anteromedial aspect and assigns the posterolateral frequency gradient to the higher-level “belt” region. Under the developmental framework that earlier stages of a sensory processing hierarchy mature before later stages, orthogonal and classical models of auditory cortex make mutually exclusive predictions for the development of the posterolateral frequency gradient. The map hA1, for example, is the most heavily myelinated tonotopic map in the region^55,29^ and should show the least developmental effect sizes compared to secondary maps. The posterolateral frequency gradient is either hA1 and shows little to no development compared to anteromedial cortex, or it is part of the belt region and should show more protracted development compared to anteromedial cortex.

To this end, we parcellated auditory cortex according to the classical framework. We identified the core as the anteromedial frequency gradient extending from the central low frequency representation on HG anterior into the high frequency representation (**Fig. 3c**, bottom), bounded on its medial extent by cortex with high myelin/neurite density (Methods). We defined the posterior half of the main diverging frequency gradient as the posterolateral section of the belt (**Fig. 3c**, bottom). Beginning with the core ROI, we extracted mean pRF parameters from each child and adult participant. The auditory core largely demonstrated stability across development, showing no significant differences in pRF preferred frequency (t(35) = −0.94, p = 0.35; **Fig. 5a**), bandwidth tuning (t(35) = 0.78, p = 0.44; **Fig. 5b**), and despite a group effect in bandwidth tuning as a function of frequency (ANOVA age x frequency interaction: F(1, 138) = 4.35, p = 0.04; **Fig. 5c**), there is no significant developmental change in the cochleotopic coverage in the auditory core (F(14, 1176) = 1.8, p = 0.16; **Fig. 5d**). Given that this classically-defined core corresponded to the anterior half of the HG ROI discussed previously, these findings suggest that the majority of HG development is driven by the posterior half of the region.

**Figure 5:**
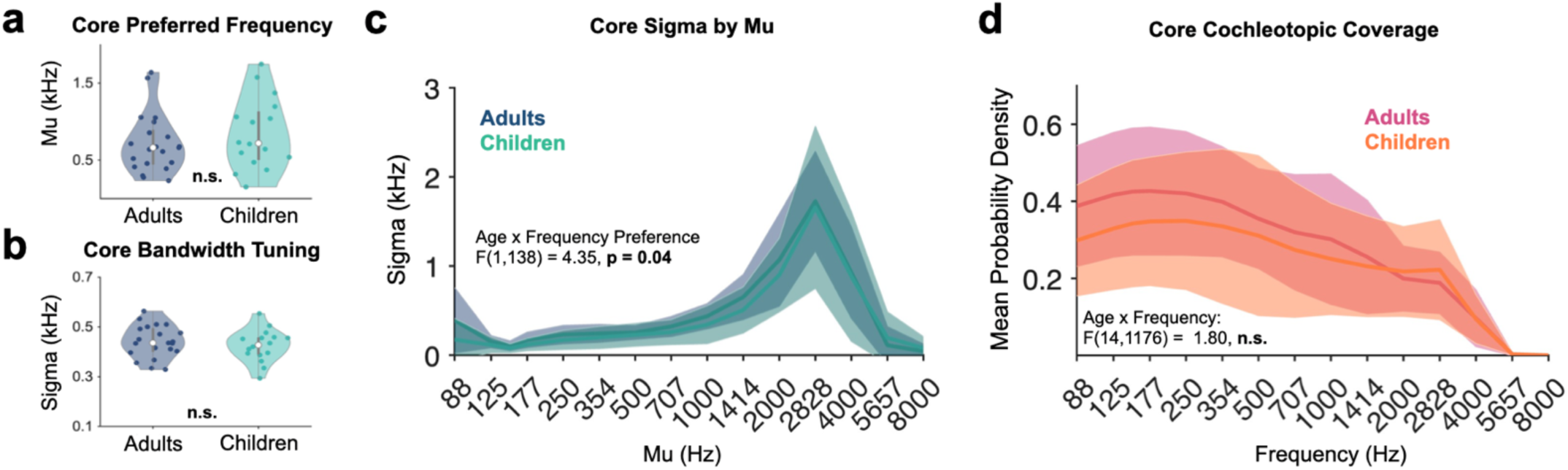
pRF properties and cochleotopic coverage of the classically-defined core are stable across development. **(a)** The mean preferred frequency (μ) across pRFs within individual participants within the core ROI. Group means denoted with white circles and standard deviation denoted with gray vertical lines. No developmental differences were observed (t(35) = −0.94, p = 0.35). **(b)** Similar to the previous panel for bandwidth tuning (σ) in children and adults. Group means denoted with white circles and standard deviation denoted with gray vertical lines. No developmental differences were observed (t(35) = 0.78, p = 0.44). **(c)** Histogram depicting mean pRF bandwidth as a function of frequency bin, separately plotted for children and adults, with shaded error bars denoting standard deviation across participants within a group. Adults and children demonstrated significantly different cochleotopic coverage patterns: ANOVA interaction of age and frequency, F(1, 141) = 4.35, p = 0.04. **(d)** Histograms showing average pRF coverage values across frequency space, solid lines denote group mean values and shaded regions denote standard deviation across participants within a group. Cochleotopic coverage pattern does not vary between groups: ANOVA interaction of age and frequency, F(14, 1176) = 1.8, p = 0.61.

Indeed, when examining the posterolateral belt ROI, we found evidence for significantly protracted pRF development. Adults showed significantly lower preferred frequency (μ) compared to children (t(35) = −2.70, p = 0.01; **Fig. 6a**), while bandwidth tuning (σ) was comparable between age groups at a mean level (t(35) = −0.75, p = 0.46; **Fig. 6b**) and across different frequencies (two-way repeated-measures ANOVA, age x frequency preference interaction: F(1, 135) = 0.22, p = 0.64; **Fig. 6c**). Comparison of cochleotopic coverage, however, revealed greater coverage in adults over low frequencies (two-way repeated-measures ANOVA, age x frequency interaction: F(14, 1176) = 11.52, p < 0.001; **Fig. 6d**). The greatest difference in coverage was observed at the lower end of frequency space, visible when visualized in a rendition of the cochlea (**Fig. 6e**), with adults having more pRF coverage compared to children at low frequencies. Children conversely showed greater average pRF coverage at higher frequencies. When examining the pRF gain parameter, while children showed larger average gain across pRFs within the core (two-sample t-test: t(35) = 3.26, p < 0.01), they showed no difference from adults within the posterolateral belt (t(35) = −1.88, p = 0.07; **Fig. S1**), and gain did not vary as a function of frequency and age (two-way ANOVA interaction: F(1, 141) = 1.17, p = 0.28).

**Figure 6:**
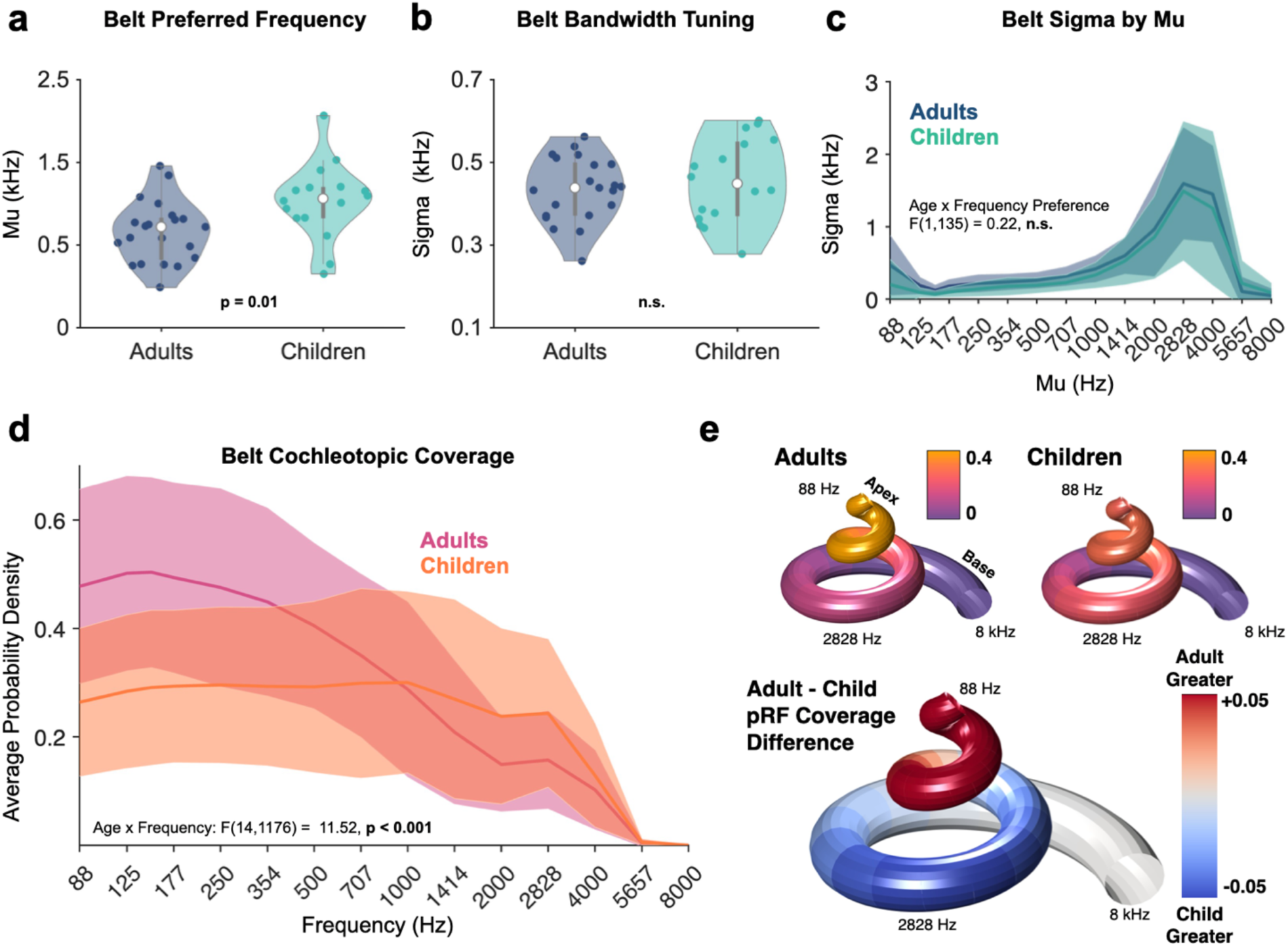
Protracted development of pRF and cochleotopic coverage properties in the posterolateral belt of auditory cortex. **(a)** The mean preferred frequency (μ) across pRFs within individual participants within the posterolateral belt ROI. Group means denoted with white circles and standard deviation denoted with gray vertical lines. Adults showed significantly lower average μ in the belt compared to children (t(35) = −2.70, p = 0.01) **(b)** Similar to the previous panel for bandwidth tuning (σ) in children and adults. Group means denoted with white circles and standard deviation denoted with gray vertical lines. No developmental differences were observed (t(35) = −0.75, p = 0.46). **(c)** Histogram depicting mean pRF bandwidth as a function of frequency bin, separately plotted for children and adults, with shaded error bars denoting standard deviation across participants within a group. Bandwidth did not vary as a function of frequency and age: ANOVA age x frequency interaction, F(1, 135) = 0.22, p = 0.64. **(d)** Histograms showing average pRF coverage values across frequency space, solid lines denoting group mean values and shaded regions denoting standard deviation across participants within a group. Coverage varied significantly across frequencies as a function of age: ANOVA age x frequency interaction, F(14, 1176) = 11.52, p < 0.001. **(e)** Visualization of cochleotopic coverage in adults (top left) and children (top right) on a model of the human cochlea. The difference of pRF coverage between adults and children (bottom) shows that adults demonstrate significantly more cochleotopic coverage than children at low frequencies while children have more coverage at higher frequencies.

### Auditory cortex structure-function coupling and development extend beyond the core and belt

With the posterolateral belt showing more protracted development in its cochleotopic coverage of auditory space than the more medially located core, these data suggested that the classical model of auditory cortex organization is more parsimonious with established developmental patterns in sensory systems. While these data also imply that there should be measurable, protracted development beyond the auditory belt, how to define functionally-distinct zones that comprise the parabelt in human auditory cortex is not yet clear. To break ground in this domain, we extended the principle of structure-function coupling to investigate tonotopy and its development in higher-level auditory cortex.

The folding of Heschl’s Gyrus in humans is variable, presenting as either a single, partially duplicated, or fully duplicated gyrus^63^. While this variability would disrupt the correlation between cortical curvature and frequency preference, the orientation of the primary high-low-high frequency gradient nonetheless follows the central orientation of HG^4^. Recent work in visual cortex suggests that this coupling between structure and function may extend beyond low-level cortical processing centers and may be a universal principle across cortex^18,66,67^. Superior temporal cortex, for example, is comprised of the sylvian fissure and the superior temporal sulcus. Emanating from the walls of the intervening superior temporal gyrus are zipper-like protrusions in the sulcal walls of the sylvian and superior temporal sulci (**Fig. 7a**).

**Figure 7:**
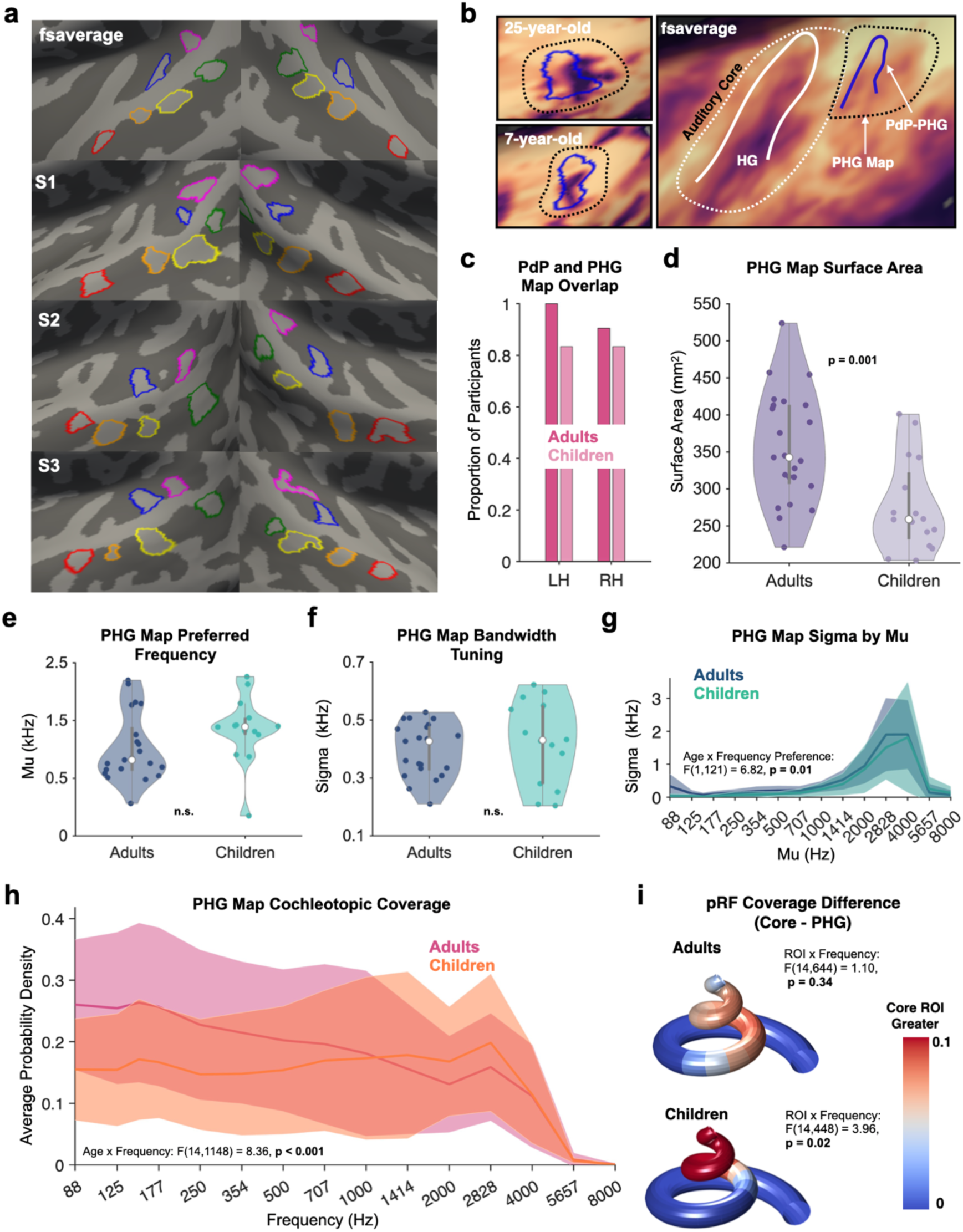
Structure-function coupling and pRF development extend beyond the auditory core. **(a)** Six plis de passage (outlined in red, orange, yellow, green, magenta and blue) were identified within auditory cortex and were consistently observed across participants, indicated by their presence on the fsaverage surface. **(b)** A PdP just posterior to HG (outlined in blue) in one example adult and one example child, which we call the PdP-PHG. The PdP overlapped with a high-low-high frequency gradient in both age groups (dotted black line). **(c)** The PdP-PHG frequently overlapped with the novel map across both age groups and hemispheres. **(d)** The surface area of the novel tonotopic map was significantly greater in adults than in children (two-sample t-test: t(37) = 3.48, p = 0.001). **(e**) The mean preferred frequency (μ) across pRFs within individual participants within PHG map. Group means denoted with white circles and standard deviation denoted with gray vertical lines. No significant developmental differences in μ (two-sample t-test: t(33) = −1.47, p = 0.15). **(f)** nor in σ (two-sample t-test: t(33) = −0.73, p = 0.47) were observed in the novel map. **(g)** Histogram depicting mean pRF bandwidth as a function of frequency bin, separately plotted for children and adults, with shaded error bars denoting standard deviation across participants within a group. Adults demonstrated significantly different average pRF tuning (σ) in low μ (μ < 150 Hz) voxels versus high μ (μ > 2000 Hz) voxels compared to children (two-way repeated-measures ANOVA with age and frequency preference as the between- and within-subject factors, interaction: F(1, 121) = 6.82, p = 0.01). **(h)** Histograms showing average pRF coverage values across frequency space, solid lines denoting group mean values and shaded regions denoting standard deviation across participants within a group. Adults and children showed significantly different cochleotopic coverage patterns: two-way repeated-measures ANOVA with age and frequency as the between- and within-subject factors, interaction: F(14, 1148) = 8.36, p < 0.001. **(i)** Greater cochleotopic coverage in the auditory core compared to the novel tonotopic map at low frequencies was observed in children (ANOVA ROI x frequency interaction: F(14, 448) = 3.96, p = 0.02), but not adults (F(14, 644) = 1.10, p = 0.34).

These small pleats in the sulcal walls are referred to as annectant gyri, originally termed *plis de passage* (PdPs) by the French anatomist Gratiolet in the 19th century in macaques^68^, and later described in humans^69^. They are buried deep within the larger sulci of the brain, sometimes interrupting sulcal continuity, and their presence early in development suggests a role in driving the functional organization of cortex^69^. Given recent work identifying PdPs in the postcentral sulcus where they colocalize with the hand-motor area^69^ and in the visual word form area of the fusiform gyrus, where they are associated with better reading performance^70^, we asked 1) if PdPs are consistently present in auditory cortex and 2) whether they act as landmarks of functional organization beyond primary auditory cortex. Identifying anatomical markers of functional topography will provide a means by which to consistently define tonotopic regions across children and adults in higher-level auditory cortex.

From structural MRI collected during the scan, we reconstructed the cortical surfaces of 21 adults and 18 children. Visualization of the inflated cortical surface revealed six small gyral protrusions–PdPs–across auditory cortex buried in the superior temporal and Heschl’s sulci, otherwise hidden in the pial view (**Fig. 7a**). We consistently observed a *pli de passage* (PdP) (**Fig. 7a and 7b**, blue outline) just posterior to HG in both hemispheres across both age groups. We referred to this novel structure by its location posterior to Heschl’s Gyrus: PdP-PHG. It is highlighted in additional participants in Supplementary **Fig. S2**. Its location just posterior to the belt ROI would place it within a cortical region consistent with the parabelt. We asked if the PdP-PHG acts as a neuroanatomical landmark that spatially co-localizes with a frequency gradient similar to HG. Across adults and children, we consistently observed a high-low-high mirror-symmetric gradient just posterior to the similarly organized tonotopy of the auditory core and belt (**Fig. 7b**, black dotted line), which we referred to as the PHG map. We defined this map’s anterior and posterior boundaries as the ridges of the high frequency representations which straddled the central low frequency representation. The lateral and medial extents of the map were dictated by the extent of the low frequency representation. This candidate map could be defined in 96% of adults and 89.5% of children and appeared consistent with a low frequency representation described in earlier tonotopic mapping work in adults^2^.

While we make no claims about the homology of this PHG map to those previously described in non-human primates^20,23,27,30^,observing spatial co-localization of this frequency gradient with a small cortical fold would afford the examination of pRF development in higher-level auditory cortex in a consistent manner across children and adults. We quantified the co-localization of the PdP-PHG and the frequency gradient with a straightforward approach, asking if the frequency gradient and the PdP overlapped consistently across brains (Boolean operation scoring 1 for a hit of any shared vertices, 0 for no overlap). In the left hemisphere, the PdP-PHG overlapped with the tonotopic map in 100% of adults and in 83.33% of children, and in the right hemisphere it was observed in 90.48% of adults and in 83.33% of children (**Fig. 7c**). We also note that the PdP posterior to HG is so consistent in its overlap with this frequency gradient that the two visibly overlap on the average cortical surface (**Fig. 7b**, right). While the high-low-high topology of this PHG map was qualitatively consistent across children and adults, we find that this tonotopic representation significantly increases in size with development (two-sample t-test: t(37) = 3.48, p = 0.001), growing from a mean of 276.68 mm^2^ in children to 356.03 mm^2^ in adults (**Fig. 7d**). This group difference persists when normalizing surface area by overall cortical surface area (Adult mean normalized area = 0.37% cortical surface area, Child mean normalized area = 0.28%; t(37) = 3.92, p < 0.001).

As in the previous ROIs, we investigated pRF properties within this PdP-PHG tonotopic map across age groups. When averaging over all pRFs within this region per participant, we observed no developmental differences in mean preferred frequency (two-sample t-test: t(33) = −1.47, p = 0.15; **Fig. 7e**) nor in bandwidth tuning (t(33) = −0.73, p = 0.47; **Fig. 7f)**. To investigate whether developmental differences in bandwidth tuning were related to preferred frequency, we compared average bandwidth tuning in low (μ < 150 Hz) and high (μ > 2000 Hz) frequency preferring pRFs between age groups (**Fig. 7g**). We found that adults had significantly larger bandwidth tuning than children specifically in low frequency preferring voxels (two-way repeated-measures ANOVA with frequency preference and age as the within- and between-subject factors respectively, age x frequency preference interaction: F(1, 121) = 6.82, p = 0.01).

To determine developmental differences in cochleotopic coverage of the PHG map, cochleotopic coverage plots were produced as described previously (**Fig. 7h**). The cochleotopic coverage curves in this region were qualitatively comparable to the auditory belt and HG ROIs: adults showed more coverage of low compared to high frequencies while children showed a flatter coverage distribution. We similarly found evidence for pRF coverage development here. A two-way repeated-measures ANOVA with frequency and age as the within- and between-subject factors respectively revealed an age x frequency interaction: F(14, 1148) = 8.36, p < 0.001, with adults showing significantly greater pRF coverage than children at frequencies ranging from 88 to 354 Hz. The coverage patterns are so distinct between groups that the peak cochleotopic coverage in adults is located between 125–177 Hz, while in children the peak is at 2828 Hz.

We next compared cochleotopic coverage differences between the classically-defined core and the PHG map within adults and children respectively. In adults, the cochleotopic coverage patterns between the core and the PHG map were not statistically different (two-way ROI x frequency repeated measures ANOVA, interaction: F(14, 644) = 1.10, p = 0.34; **Fig. 7i**, top). Children demonstrated significantly different coverage patterns between their core and PHG maps, with attenuated coverage of low frequencies in the PHG map compared to the core (ROI x frequency interaction: F(14, 448) = 3.96, p = 0.02; **Fig. 7i**, bottom). Thus, the developmental increase in PHG map coverage of low frequency space demonstrates that this region of auditory cortex has not yet reached its characteristic, adult-like patterns of cochleotopic coverage by middle childhood.

### Auditory pRF development is behaviorally relevant

Because the participants who underwent neuroimaging also completed psychoacoustic tests (**Fig. 1a**) outside of the scanner, we asked whether the developmental differences in pRF properties are behaviorally relevant. To this end, we investigated the relationship between frequency discrimination thresholds at 150 Hz, 2000 Hz and 4000 Hz–as measured in the 3-AFC tone-in-noise task–and cochleotopic coverage at these same frequencies. Since we have no prior for whether low versus high-level pRF computations would be the most behaviorally relevant, we chose to use pRF coverage from the broader HG ROI given that it 1) is inclusive of both the core and posterolateral belt, and 2) inclusive of both organizational frameworks. Further, we focus on coverage as a neural metric given the significant developmental differences in HG cochleotopic coverage (**Fig. 4d, 6d**), and the fact that coverage inherently incorporates both μ and σ from a region’s pRFs and is thus a more holistic metric.

To determine how age and frequency of the tone may interact with cochleotopic coverage to affect frequency discrimination thresholds while accounting for interactions with age and frequency, we fit a linear mixed-effects model (LME) to the data, with age, frequency and cochleotopic coverage as fixed effects and individual participant slopes as the random effect (**Fig. 8a-8b**). We found a significant interaction between the factors of age, frequency, and coverage: F(2, 220) = 5.05, p = 0.007. A Holm-corrected post hoc test revealed this effect to be significant at 150 Hz (F(1, 74) = 8.76, p = 0.01). This interaction of frequency, coverage, and age suggests that as pRF coverage increases for a given frequency, the detection threshold likewise improves, and this relationship is stronger in adults than children. While there are changes in the coverage and threshold associated with each frequency, this effect is being driven predominantly by changes in pRF coverage and detection of the 150 Hz tone.

**Figure 8:**
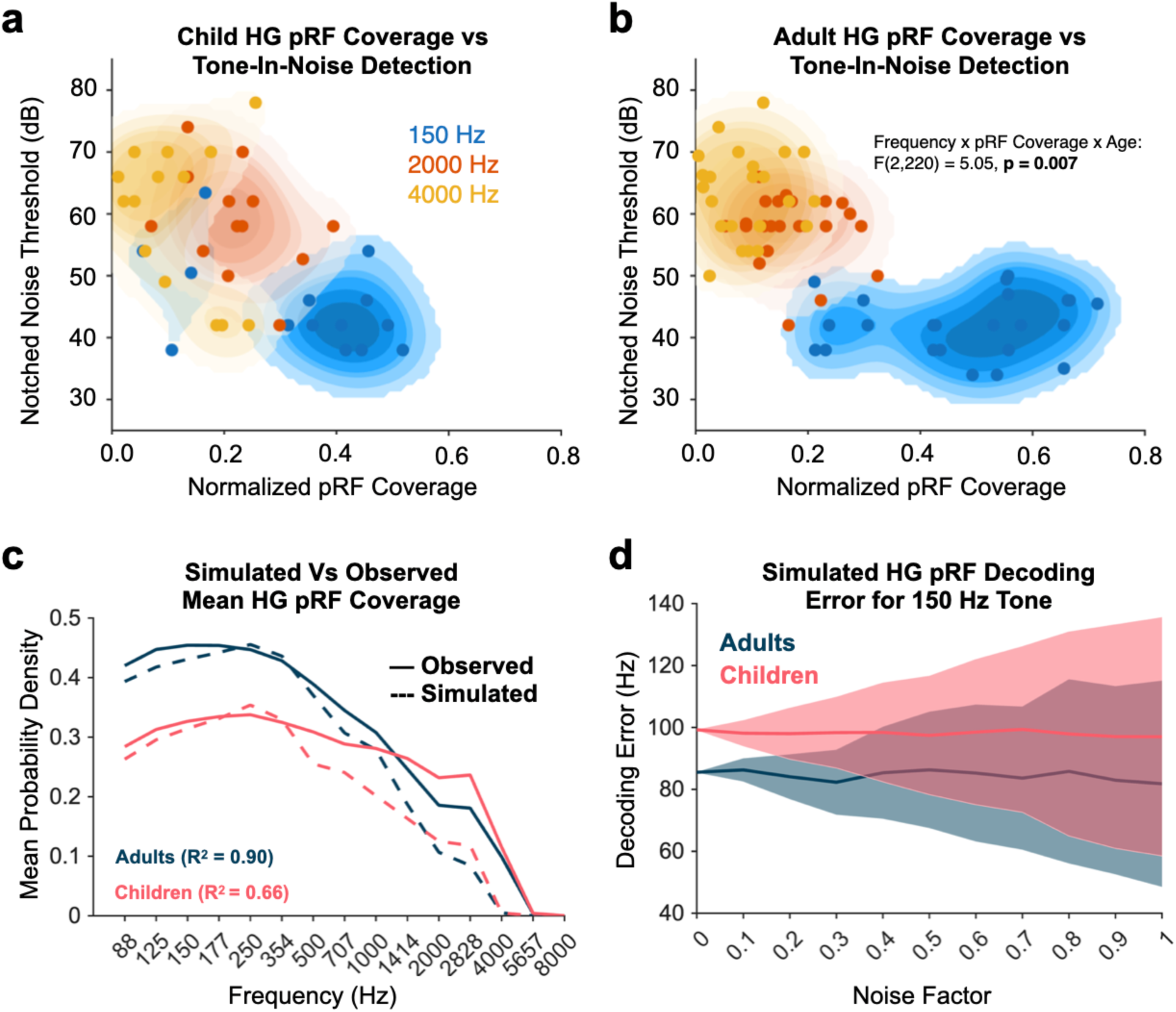
Auditory pRF development is behaviorally relevant. **(a)** A linear mixed-effects model determining the relationship between cochleotopic coverage in auditory cortex surrounding HG (orthogonal hR and hA1, or classical core and posterolateral belt) and tone-in-notched noise detection thresholds in children. Age, frequency and cochleotopic coverage were fixed effects and individual participant slopes were modeled as a random effect. **(b)** Similar to the previous panel but for adult data. The linear mixed-effects model revealed a significant interaction between all three factors: F(2, 220) = 5.05, p = 0.007. **(c)** The cochleotopic coverage of the simulated receptive field populations produced in adults and children overlaid with the observed coverage patterns. **(d)** Decoding error resulting from the stimulus reconstruction simulation in children and adults. Central solid lines denote group means, and shaded regions denote standard deviation. While decoding error variability increases with additional noise, larger bandwidth tuning and heavier pRF coverage of low frequencies confers less decoding error across noise levels in adults compared to children.

Previous work in visual cortex demonstrated that increasing pRF coverage of a given point in visual space, whether through spatial attention allocation^71^ or developmental increases^18^ in pRF coverage, reduces the uncertainty of a given visual stimulus’ location. We thus investigated how our observed developmental increase in pRF coverage over low frequencies may aid in the perception of these stimuli. To that end, we simulated populations of adult and child pRFs derived from the mean pRF parameters from the broader HG ROI in each age group respectively (Methods). Because we observed broader bandwidth tuning (σ) in adults than in children specifically in low frequency (μ) voxels, we assigned the mean σ in low μ voxels (278 Hz in adults and 210 Hz in children) to all simulated voxels of the adult and pediatric population, respectively. Preferred frequency (μ) values were assigned to each simulated voxel to produce the best reconstruction (based on least-squares regression) of the mean pRF coverage plot for each age group (**Fig. 8c**). The decodability of a frequency was determined by how well the response of a population of pRFs could reconstruct the stimulus, measured by the difference (in Hz) between the predicted and actual tone. To more closely relate this simulated decoding task to our experimental tone-in-noise detection task, we assessed how decoding ability is affected by noise. Specifically, we added Gaussian white noise to the population response, with a standard deviation representing a proportion of the mean population response (1-100%).

For each additive noise factor, decoding error was assessed across 1000 iterations and subsequently averaged across the iterations (**Fig. 8d**). A linear mixed-effects model with age and noise as the fixed effects and iterations of the simulation as the random effect found that the adult pRF population produced a lower decoding error for the reconstruction of a 150 Hz tone compared to the child pRF population (main effect of age: F(1, 21996) = 614.08, p < 0.001) and that this was consistent across increasing noise levels (age x noise interaction: F(1, 21996) = 0.23, p = 0.63).

## Discussion

Our results show that the tonotopic maps which occupy the cortex of Heschl’s Gyrus, such as the anteromedial core and posterolateral belt, are qualitatively present by middle childhood. However, investigation of auditory pRF properties revealed that neural populations undergo protracted development, more densely sampling the lower end of frequency space in adults compared to children. Given that this functional maturation was most dramatic in the posterolateral aspect of HG, compared to the developmental stability of pRF properties in the anteromedial section, these data offer evidence for a form of hierarchical development consistent with an organizational framework which places the primary “core” of auditory regions on the anterior half of HG and the secondary “belt” on the posterior half. Importantly, this neural development was behaviorally relevant. We observed better tone-in-noise detection thresholds in adults compared to children specifically for a low frequency which was related to individual differences in pRF coverage at that same frequency. Lastly, we identified the presence of a novel *pli de passage* in both age groups which consistently overlapped a mirror-symmetric tonotopic gradient outside of the auditory core and belt. Having overcome hurdles to pediatric pRF mapping, these results offer a portrait of the protracted development of human auditory cortex: tonotopic representations in higher-level auditory cortex more heavily sample low frequencies into adulthood which mirrors behavioral improvements in low frequency sensitivity.

Using a novel tonotopic mapping procedure adapted for children, we find that tonotopic organization in auditory cortex is qualitatively stable by middle childhood (**Fig. 3a, 3b**), with adults and children showing a similar spatial arrangement of a high-low-high frequency gradient across HG. The congruent presentation of auditory (tones) and visual (robots) stimuli may have improved SNR of auditory data given prior work showing that audiovisual stimuli enhance responsiveness in auditory cortex. Further, as previous studies have demonstrated that visual speech can influence auditory cortex activity^72–76^, it is possible the perception of robots emitting tones could impact BOLD signals. However, the cartoon robots of *RoboTones* lacked any facial animation, reducing the perception of visual speech. Still, the moving robots may have elicited the perception of auditory stimulus motion, potentially activating the planum temporale which has shown to be involved in auditory motion direction^77^. Nonetheless, any such effects would be present in both age groups and should not impair interpretation of developmental effects. The gamified pRF mapping approach allowed us to identify and quantify the development of auditory receptive fields across multiple stages of the auditory processing hierarchy.

To date, auditory pRF mapping studies have offered competing frameworks for how to map tonotopic fields established in non-human primate work onto the superior temporal plane of humans. For example, the orthogonal and classical frameworks differ in the localization of the primary core relative to HG: the orthogonal model places core maps hR and hA1 across HG, whereas the classical interpretation places the core along the medial aspect of HG, parallel with its long axis. The classical framework offers a compelling macaque-like orientation of the core that is consistent with cytoarchitecture (i.e. a well-developed layer IV^27^) as well as accounts of a heavily myelinated primary sensory region A1 in the medial aspect of HG^29,55^. However, the orthogonal view recognizes the evolutionary expansion of the superior temporal plane which resulted in a HG that can be observed in humans but not most non-human primates^78^, complicating cross-species comparisons of functional topography. While development could be observed in the orthogonal core, defined as a broader ROI encompassing both sides of HG (**Fig. 4**), parcellating HG into the anteromedial “core” and the posterolateral “belt” revealed that this development was not homogeneous and instead driven by protracted development in the posterolateral belt (**Fig. 6**). Importantly, this development was not observed in the classically-defined core–consistent with work in the visual domain showing adult-like pRFs in early visual areas but a protracted development of pRF properties in higher-level regions^19^. Given that the orthogonal model places hA1 where the classical model places the belt, the classical model of auditory cortex is more parsimonious with developmental frameworks of cortical sensory system development in the human brain^48^.

Our comparison of pRF properties in auditory cortex between age groups shows protracted development in the way auditory neural populations sample frequency space. With age, pRFs in HG and the posterolateral belt more densely sample the lower end of frequency space. This is consistent with prior work showing that air-cell volume of the mastoid process of the temporal bone increases with development^37^ and is theorized to promote better transduction of low frequency sounds^35^. This work also provides a neural correlate for tympanometric mapping work in children which suggested a developmental improvement in pure tone detection primarily for lower frequencies^33^. Our work extends these prior findings to show that detecting tones embedded in noise also undergoes protracted development, especially in very low frequencies such as 150 Hz, which was not tested in prior work^35^. When simulating how increases in pRF coverage interact with population-coding of stimulus identity, we find that the greater coverage of low frequency space in adults provides a stimulus identity representation that is less noisy and more robust to external noise compared to children (**Fig. 8c-d**). This is in line with work on visual pRFs which reported that visual attention similarly serves to increase pRF coverage and spatial certainty of a stimulus. While prior work on modeling attentional effects on auditory pRFs finds sharpening of pRF tuning width^79^, concurrent shifts in preferred frequency may improve population sampling of the attended frequency to further aid perception. Indeed, some work has shown that attending to a single high frequency tone can improve responses of neural populations preferring nearby frequencies^54^, suggesting potential recruitment of a larger population to improve coverage.

In our behavioral *Noisy Nest* paradigm, it should be noted that there were differences in the size of the extent and notch of noise bands across frequencies. Prior work demonstrated that extending noise beyond a central frequency in a detection task past a certain bandwidth has no impact on a listener’s ability to detect the central tone^61^. Importantly, work mapping the size of this noise bandwidth demonstrated that while this measure scales with frequency, the noise bandwidth for detecting a 10 kHz tone is only around 1000 Hz wide^80^. Because all of the noise masker bandwidths in our paradigm extend 2500 to 4000 Hz beyond the central frequencies, there should be no differential impact on detection thresholds across frequencies. Differences in notch size within the noise masker can impact the detection of a central tone depending on how much the noise overlaps with the “auditory filter” for the target frequency^35,61^. Relative to the other frequencies, the notch for the 150 Hz tone is relatively wider, which would make detection of this tone easier than average. While this can impact our ability to infer behavioral differences between frequencies, it is still the case that children demonstrate higher thresholds for detecting this tone compared to adults (**Fig. 1e**) given that experimental parameters are consistent across groups. The fact that this condition is relatively easier than others suggests that the developmental difference we report here is likely a conservative estimate.

Examining cortex outside of the auditory core and belt, we find evidence that higher-level regions of auditory cortex undergo protracted development. Here, we characterize a tonotopic map posterior to the lateral belt ROI in both children and adults (**Fig. 7**). While in adults this map shows relatively greater sampling of low compared to high frequency space, it shows a flatter coverage distribution in children (**Fig. 7h**) which significantly differs from the coverage pattern in the core (**Fig. 7i**). This map also appears to be consistently yoked to an anatomical feature known as a *pli de passage*, a small gyral ridge branching from the superior temporal gyrus into the sulcal bed posterior to HG. Our characterization of this previously undefined *pli de passage* reveals that it was consistently present in both adult and child participants (**Fig. 7c**). Prior work in adults^2,29,32^ visualizing function and structure together reported a low frequency representation posterior to Heschl’s Gyrus, although this region has not been explicitly linked to underlying anatomy. Our finding of an additional tonotopic mirror-reversal posterior to HG, one that co-localizes with a small cortical fold, replicates this earlier work and provides further evidence for consistent and anatomically-grounded tonotopic maps beyond the core and belt in human auditory cortex.

Due to high inter-subject variability in the morphology of the superior temporal plane, the relative orientation of the PdP in relation to the novel tonotopic gradient varied across participants but consistently overlapped with the tonotopic map in more than 80% of participants (**Fig. 7c**). The consistent proximity between this PdP and the tonotopic map suggests a strong link between structure and function which recapitulates similar observations in the visual domain^18^. A PdP may be the cortical fingerprint of U-shaped white matter fibers that interconnect neural populations on distal sides of sulcal walls^81^, and may thus capture distinct connectivity zones. Because the PdP-PHG overlaps with a zone representing a complete high-low-high frequency representation, the PdP may serve to accommodate axonal input from its likely afferents originating from the primary frequency gradient on HG.

This coupling between structure and function also allowed us to consistently define an auditory region outside of the auditory core and belt–for which there is currently no standard procedure in humans–to investigate its development. We found that this tonotopic map exhibits a cochleotopic coverage pattern distinct from that of the auditory core in children (**Fig. 7i**) and undergoes protracted developmental increases in its sampling of low frequency space (**Fig. 7h**). Because the medial-lateral extent of this map is defined according to the extent of its central low frequency representation, it may be the differences in low frequency pRF tuning between groups which drive the difference in size of this map (**Fig. 7d**), which suggests the group-differences we observe in cochleotopic coverage may be a conservative estimate. That is, if the true map borders in children are larger, they would extend further into the surrounding high frequency representation and further reduce the relative representation of low frequencies reported in **Figure 7h**. Further, prior work has implicated the planum temporale in auditory motion direction perception^77^, suggesting its role in higher-order auditory computations. Our finding of protracted development in a tonotopic map which overlaps with this region is thus in support of hierarchical models of brain development^18,19,47,48^ in which higher-level regions of the sensory processing hierarchy show the most prolonged development.

Importantly, the tonotopic map we describe overlapping the posterior PdP appears to be a continuation of the high-low-high frequency gradient of HG (**Fig. 7b**), as previously noted in adults^2^. Thus, the reversal of the frequency gradient from the core into the surrounding auditory cortex mirrors retinotopic organization of visual field maps, whose borders occur at polar angle reversals^82^. Because PdPs are present early in development^69^ and the location of sulcal landmarks relative to HG as well as the number of HG duplications are similar in adults and children^63,83^, we hypothesize that cortical folds act as early scaffolds for functional organization later in life. Future work should investigate if the remaining *plis de passage* we described here extending from the STG into the STS and superior temporal plane also landmark other tonotopic representations. If so, PdPs may represent a principled, neuroanatomical metric by which to parcellate the functional organization of human auditory cortex more consistently across individuals.

One possibility worth considering is that the development of pRF tuning observed across auditory cortex is driven by auditory experience. While the general bias towards greater coverage of low (<1000 Hz) frequencies is evident across all participants (**Fig. 4d**) and may be driven by logarithmic sampling techniques of pRF mapping, adults demonstrated more pRF coverage for low frequencies than children for frequencies 88-350 Hz which, notably, overlaps with the frequency range of the human voice which covers approximately 88-290 Hz across male and female speakers^84^. Results from our behavioral experiment show that tone-in-noise detection performance is still developing in children at 150 Hz which also falls within this frequency range. Given that high-level auditory cortex shows greater selectivity for human voices with age^83^ and that voice recognition performance improves with age^59^, it is possible that pRFs are re-tuning with development to more densely sample from an ecologically relevant frequency range in support of human voice processing. Indeed, the spatial co-localization of voice-selective regions with cortex tuned to lower frequencies^85^ supports the relationship between voice processing and tonotopic organization. This would be consistent with findings in the visual domain where face-selective pRFs become more foveally biased with age, presumably to support the processing of faces which tend to fall within the center of the visual field^19^.

Regarding our tone-in-noise behavioral paradigm, previous studies have used this approach to estimate the width of the parallel bank of bandpass filters in the peripheral auditory system^35,36^. Perceptually, the width of a given filter corresponds to the certainty that a frequency is present amidst noise, with a narrower filter being less prone to surrounding noise. Prior work^35,36^ has demonstrated that the auditory filter potentially narrows from the age of 6 years old to adulthood. While it is certainly the case that narrower tuning of an individual neuron could support improved selectivity for a given frequency, it should be noted that estimates of the auditory filter width are based on the perception of behaving humans, and it is therefore likely this effect emerges from the broader population of cells engaged in audition from cochlea to cortex. We can draw a parallel with receptive field mapping work in visual cortex which shows that spatial attention serves to improve pRF coverage density at a particular point in space which in turn reduces uncertainty about the spatial position of a visual stimulus^71^. This effect has also been observed in developmental contexts, where maturing pRF properties in visual field maps serve to improve coverage of a given point in visual space and improve the spatial coding of a visual stimulus^18^. In our data, we hypothesized that the developmental improvement of pRF coverage at low frequencies (**Fig. 4d**) may serve to improve the certainty of perceiving low frequency sounds. Indeed, the results of our frequency decoding analysis revealed that an adult-like pRF coverage pattern resulted in the improved decoding of a 150 Hz tone compared to its child-like counterpart.

One limitation of this study is that while we mapped pRF tuning across frequency space, it is known in auditory cortex that periodicity of sounds is represented orthogonally to frequency^86^. Thus, the organization and development described here is only one dimension of relevant auditory computations. Second, we delineated the tonotopic map boundaries of our auditory ROIs using the same pRF mapping data that was used to estimate pRF properties. Though this is a common practice in pRF mapping of visual cortex^57,82^, future work can utilize probabilistic maps based on cytoarchitecture or anatomy to define ROIs independently of their function. We do note that there was no significant difference in the size of the auditory core between children and adults (**Fig. 3d**), suggesting that our definition of the core was not biased in any way between groups and could not have driven the differential pRF development we observed. The difference in the size between groups for the PHG map, however, could be driven by the relatively smaller amount of low frequency representation in this region in children (**Fig. 7**) given that the lateral-medial extent of the PHG map was drawn according to the low frequency representation at its center. In this case, the developmental difference we report here for the PHG map (**Fig. 7h**) is likely a conservative estimate; extending the map along the medial-lateral axis to the adult-like size would include even more high frequency preferring pRFs and further distinguish the age groups.

One potential source of noise was the presence of scanner noise during our pRF scanning procedure which other labs have mitigated using a sparse imaging protocol^87^. Given time constraints in pediatric neuroimaging and the desire to maximize voxelwise SNR, we opted instead for a standard fMRI protocol and employed in-ear microphone tips that reduce scanner noise by 32 dB while simultaneously delivering auditory stimulation. The matched pRF model fits (**Fig. 2c, 2d**) and qualitatively similar size and topology of the auditory core and belt (**Fig. 3d-3f**) in children and adults suggest that this approach was equally effective in both groups. A further reassurance that scanner noise was not introducing developmental artefacts into pRF data comes from our out-of-scanner behavioral data. Children show poorer detection thresholds only for low frequencies, mirroring the neural data measured independently in the scanner. Lastly, both children and adults show a general relationship between pRF coverage and tone-in-noise detection of the three different frequencies, suggesting a link between the data measured inside and outside of the scanner.

## Methods

### Participants

We collected neuroimaging data from 25 children ages 5-11 years (mean age 9.00 ± 1.63 years, 12 females) and 25 adults ages 21-35 (mean age 25.72 ± 2.82 years, 16 females). The age range of children was selected to capture the developmental trajectory of both low- and high-level auditory object processing as established by previous work^33,59^. Our adult age range was chosen to match that of our previous developmental work^19^. Data from 5 children were excluded due to excessive head motion, and data from 2 other children and 4 adults were excluded after matching age groups for variance explained by the pRF model in our comprehensive auditory ROI. We report the pRF mapping results from the remaining 18 children and 21 adults. All participants were recruited with normal hearing and were approved by the Institutional Review Board of Princeton University. Prior to their participation, adult participants and parents of child participants provided written informed consent, and children provided written assent. Adults and children received $30 per hour for scanning and $20 per hour for behavior, and children additionally received a small toy after each session for their participation.

Neuroimaging data was collected over two sessions in children. At the beginning of the first MRI session, children were trained in a simulated scanner to acclimate them to the MRI environment. During the training, they watched a 5-minute movie in this mock scanner while receiving biometric feedback to help minimize head motion. Children who were able to successfully lie still (less than 2.4 mm of head motion) for the duration of the training were invited to participate in the rest of the session. We next collected structural MRI from children which was used to produce cortical reconstructions of the brain. Lastly, children completed a behavioral task outside of the MRI machine. Children returned for the second session to complete functional MRI experiments within two months of the initial visit. During this session, participants completed the *RoboTones* auditory pRF mapping experiment. Adults underwent structural and functional MRI and completed the behavioral task in one session.

All participants provided written and informed consent. Child participants provided assent alongside parent consent. All experiments and procedures were reviewed and approved by the Princeton University IRB, protocol 13074.

## Data Acquisition

### Structural MRI

Data were collected on a 3T Siemens Magnetom Prisma MRI scanner at the Scully Center for the Neuroscience of Mind and Behavior within the Princeton Neuroscience Institute at Princeton University using a 64-channel phased array coil. T1- and T2-weighted anatomical images were collected using a 3D MPRAGE sequence (TR = 2300 ms, TE = 2.96 ms, TI = 900 ms, flip angle = 9°, field of view = 256 × 256 mm, matrix size = 256 × 256, voxel size = 1.0 × 1.0 × 1.0 mm³).

### Functional MRI

Functional data were collected on the same scanner and head coil as the structural images. fMRI scans were acquired using a 2D echo planar imaging (EPI) sequence (TR = 2000 ms, TE = 34 ms, flip angle = 80°, voxel size = 2 × 2 × 2 mm³, 40 slices, slice thickness = 2 mm, field of view = 192 mm, matrix = 96 × 96). Each run was 400 seconds in duration and there were three runs in total. As in our prior work in visual cortex^18,19^, runs were averaged to produce a single higher-SNR run which was later used for fitting population receptive fields.

### pRF mapping experiment

During functional MRI, adults and children completed three runs of an auditory pRF mapping experiment in which they were presented with pure tone frequencies. The experiment was adapted from previous work^57^ to be more compatible with pediatric neuroimaging. The following changes were made: 1) three 7-minute scans were collected instead of the original six 9-minute scans 2) runs were averaged to improve SNR, and 3) the task was gamified to better engage younger participants as described below. Before starting the game, titled *RoboTones*, participants were given task instructions and had the opportunity to practice outside of the scanner. In the game, the player works in a robot making factory and must determine which robots are broken and need to be sent back for repair. A series of cartoon robots move horizontally across the screen on a conveyor belt and each “beeps” a pure tone frequency. Participants responded with a button press when they heard a robot repeat its tone two times in a row, indicating that it was broken. Each 400 second run consisted of 10 blocks–in each block, frequencies ranging from 88 to 4000 Hz in half octave intervals (88, 125, 177, 250, 354, 500, 707, 1000, 1414, 2000, 2828 and 4000 Hz) were presented for 2.5 seconds each in semi-random ascending and descending sweeps. Tones were presented in a random Morse code-like pattern to increase saliency over scanner-induced noise as well as to activate both “on” and “off” auditory populations. Auditory stimuli were delivered via an MRI compatible audio delivery system (Avotec Silent Scan SS-3300) at a sampling rate of 44.1 kHz. Sound intensities were equalized for perceived loudness according to the ISO 226 equal loudness curve^88^. Sound levels were not adjusted in individual participants such that any differences observed in tonotopic organization between the groups reflected developmental differences and not differences in auditory stimulation. Three identical runs were collected from each participant to be averaged across and therefore increase SNR.

### Behavior

Outside of the scanner, 32 children ages 5-11 years (mean age 8.47 ± 1.61 years, 17 females) and 35 adults ages 21-34 years (mean age 26.06 ± 3.32 years, 18 females) completed a gamified 3-alternative forced choice (3-AFC) tone-in-noise detection task adapted from a previous study^35^. Before starting the game, participants were instructed on the task: their goal was to help a mama bird determine which of her eggs was ready to hatch based on the sound it made. In the game, participants were presented with 3 colorful eggs which, one-by-one, shook and produced a sound for a 2 second duration with an ISI of 0.5 seconds. The sound was either a pure frequency tone embedded in white noise or white noise alone. After listening to each sound, the participant had to choose the egg which emitted the target tone embedded in white noise, indicating that it was ready to hatch. Feedback was provided for correct responses as a cartoon chick hatching from the egg while the auditory feedback “No…” was given for incorrect responses. The location of the correct egg was randomized for each trial and stimulus presentation order was the same across participants. Tones were either 150 Hz, 2000 Hz or 4000 Hz pure tones and could be masked by either flat spectrum white-noise or by white noise with a notch centered at its frequency. Target tone stimuli were generated in MATLAB by determining a sine function at a given frequency over a 2 second time interval. The white noise maskers were also generated in MATLAB as 2 second long samples of Gaussian white noise. In the flat-spectrum condition, the white noise was a 4000 Hz wide band centered at the target frequency (except in the 150 Hz condition where the masker covered the frequency range 10 Hz to 2500 Hz). In the notched spectrum condition, the masker comprised two 2000 Hz wide bands separated by a notch at the signal frequency. In the 150 Hz condition, the two bands covered 10 to 50 Hz and 250 Hz to 2250 Hz, resulting in a notch width of 200 Hz. The 2000 Hz and 4000 Hz notch spectrum noise conditions had spectral widths of 800 Hz and 1600 Hz respectively. In both white noise conditions, the SPL spectrum inside the white noise band was 60 dB greater than outside the band. Auditory stimuli were presented via PsychPortAudio through over-ear headphones to reduce environmental noise during the task. The sound level of the task was kept consistent across individuals. The task consisted of 6 blocks: 1) 150 Hz notched white noise 2) 150 Hz flat spectrum white noise 3) 2000 Hz notched white noise 4) 2000 Hz flat spectrum white noise 5) 4000 Hz notched white noise 6) 4000 Hz flat spectrum white noise. To determine participants’ thresholds for perceiving each tone masked by white noise, the Quest+ staircasing procedure^60^ was used to adaptively adjust the sound level (dB) of the tone based on the correctness of the response. The likelihood function was defined as a Weibull distribution with the following parameters: slope = 2, gamma/guess rate = 0.33 (for 3-AFC), lambda/attention lapse rate = 0.05, threshold = 30 - 100 dB. A binary response domain was used (0 = “no”, 1 = “yes”), and an entropy of 3 was used as the stopping criterion for determining the threshold for detecting the masked tone after a minimum of 20 trials. The visual components of the game were generated in PsychtoolBox.

## Data Analysis

### Anatomical data analysis

T1- and T2-weighted anatomical images were analyzed using FreeSurfer version 7.1.1, resulting in gray and white matter segmentations that were used for cortical surface reconstruction and visualization of tonotopic data in each participant’s native space^89^.

### fMRI data analysis

fMRI data was preprocessed using the HCP processing pipeline for motion correction, slice-timing correction, and top-up phase distortion correction. FsFast (FreeSurfer Functional Analysis Stream; https://surfer.nmr.mgh.harvard.edu/) was used to resample voxel time-series data to the native brain surface. No spatial smoothing was applied. Participants who had a mean framewise displacement (FWD) near 1 voxel (2 mm) within a scan were excluded from further analysis or invited back to try again in a subsequent session. Though children had significantly more head motion than adults, both groups had mean FWD values below 0.5 mm as used in prior work^62^. Furthermore, age groups were matched for mean variance explained by the pRF model (**Fig. 2d**), ensuring no differences in data quality (see Results).

### pRF Modeling

From our fMRI data, we performed population receptive field (pRF) modeling, a computational approach that allows for the quantification of pRF properties in each voxel based on its BOLD response to sensory stimuli. We implemented the model fitting procedure from Thomas and colleagues^57^. pRF modeling started with defining the stimulus time course which indicates the presence or absence of an auditory stimulus in time and log frequency space. This stimulus image (**Fig. 2b**) was then convolved with the hemodynamic response function (HRF) to create a predicted time-course. The pRF of a voxel was defined as a 1-D Gaussian with frequency center corresponding to its preferred frequency (μ) and a standard deviation (σ) that approximates bandwidth tuning, all in units of the mapping stimulus, Hertz. The predicted and observed fMRI time-series of a given voxel were compared in an iterative procedure which optimized the pRF parameters that best predicted the observed time series by maximizing their correlation. After fitting, only voxels that had a variance explained by the pRF model equal to or greater than 10% and a pRF with standard deviation between 0.01-2 in log frequency space (as in previous work^57^) were included in further analyses. Software repositories for auditory pRF modeling were developed by the University of Washington Vision and Cognition Group. The pRF parameters of each voxel were used to visualize frequency representations in each participant’s native space. Group averaged tonotopic maps were produced by resampling each participant’s tonotopic map to the fsaverage surface using cortex-based alignment^89–91^ and then for each vertex, averaging pRF parameters across all participants. We ensured that pRF model fit quality was similar between groups in an independently-defined ROI encompassing the superior temporal cortex (**Fig. 2d**) which included the planum polare, HG, the planum temporale, and up to the ridge of the superior temporal gyrus.

### Myelin Mapping Using T1-w/T2-w Ratio

For each participant, the voxel-wise ratio of their T1-to T2-weighted images was calculated to generate a volumetric myelin contrast image. FreeSurfer’s mri_vol2surf function was then used to sample the volumetric contrast values onto the cortical surface within the gray-matter ribbon, using a fractional distance corresponding to the midpoint between the white-matter and pial surfaces. The resulting surface-based myelin contrast maps were restricted to a comprehensive auditory cortex ROI (see below). Within each participant and hemisphere, vertices were ranked according to myelin contrast within the auditory cortex ROI, and a label containing the top 10% of vertices was generated. To generate group-averaged myelin maps, each participant’s native-space myelin map was registered to the fsaverage cortical surface using FreeSurfer’s mri_surf2surf. The transformed maps were then averaged at each vertex across participants within each age group (n = 25 adults, n = 18 children). Within each group-averaged map, vertices were again ranked according to myelin contrast within the fsaverage auditory cortex ROI, and a label containing the top 10% of vertices was generated to define the region of highest myelination (**Fig. 3b**, bottom, solid black outline). The myelin-dense region of HG was used to define the medial boundary of the classically-defined core on the anteromedial aspect of the low-to-high frequency gradient (**Fig 3**).

### ROI Definitions

Auditory cortex regions of interest were defined according to two frameworks, described below.

Classical Model:

A region of interest (ROI) of the human auditory core was drawn in each hemisphere of each participant’s inflated cortical brain surface using drawing tools in FreeSurfer. A map was only defined after consensus was reached between the authors. The orientation of the core was drawn in correspondence with the macaque model, lying along the long axis of HG and within the anteromedial region of the gyrus. The medial portion of the core was delineated as the region of greatest myelination and its lateral edge corresponded to the base of HG (**Fig. 3c**, bottom). While this core ROI contains the subfields hA1, R, and RT which share a low frequency border, individual subfields were not defined and the core ROI was analyzed as a single structure. A lateral belt ROI was similarly drawn in accordance with the macaque model of auditory cortex. A single high-to-low tonotopic gradient running postero-medially to antero-laterally was observed in all participants, likely encompassing lateral belt subfields ML and AL. Because the high-low-high frequency gradient which characterized the human auditory medial belt in prior work^29^ was not consistently observed across our participants, we did not define a medial belt ROI.

Orthogonal Model:

An additional ROI was drawn in each participant to include all voxels within a contiguous region of auditory cortex that encompassed a mirror-symmetric high-low-high frequency gradient consistent with the “orthogonal” orientation of the core. We refer to this ROI as “HG” given that the gyrus was used as an anatomical landmark for the low frequency portion of the tonotopic gradient. Subfields hA1 and hR each comprised a low-to-high frequency gradient, extending postero-laterally and antero-medially, respectively from the low frequency portion. Anterior and posterior borders were aligned with the central ridges of the high frequency representations anterior and posterior to HG. Lateral and medial borders encompassed the entire medial-lateral axis of HG as in previous work^4^.

We also defined a separate ROI to include the auditory core and belt as well as surrounding auditory cortex in the superior temporal plane and the superior temporal gyrus in which we could assess pRF model fit between age groups independent from the auditory ROIs in which we would be analyzing pRF properties. The mean variance explained of the pRF model fit was extracted from this control ROI in each participant to assess data quality (**Fig. 2d**).

A functionally defined ROI was produced to examine pRF development in a second tonotopic map posterior to HG overlapping a *pli de passage*. While we make no claims to the homology this representation may have with auditory fields defined in non-human primates, we use this region here with the goal of quantifying functional development in (i) a region outside of the auditory core and belt, and (ii) a region matched for the high-low-high topology of primary auditory cortex. We refer to this map as PdP-PHG. This candidate tonotopic map was defined based on the presence of a mirror-symmetric high-low-high frequency gradient just posterior to the HG ROI, such that they shared a high frequency representation (**Fig. 7b)**. That is, the high frequency ridge defining the posterior extent of the HG ROI was the beginning of the PHG map and thus marked its anterior border. The posterior border of the PHG map was drawn at the ridge of the next high frequency region posterior to the PdP to produce a map with a symmetric high-low-high pattern similar to the HG ROI. Lateral and medial borders were drawn according to the medial-lateral extent of the low frequency representation.

**PdP Identification:**

PdPs were identified based on cortical morphology within our comprehensive auditory ROI. FreeSurfer was used to produce a surface of the grey/white matter interface in each participant. As in recent work^81^, PdPs were defined on this white surface as the presence of a “wall-pinch” where a sulcal wall bulges slightly to form a ridge. As observed in the STG^81^, PdPs either presented as a complete connection of the 2 sulci (superficial) or were sometimes partially buried, making this ridge hidden or only partly visible on the white surface (deep). The presence of a PdP was confirmed on the inflated cortical surface, where they presented as small gyri. Six PdPs were first identified on the fsaverage cortical surface within our anatomical ROI and were then projected onto each participant’s native cortical surface (**Fig. 7a**) to provide a neuroanatomical prior when identifying the PdPs in each participant’s individual brain. PdPs in the native cortical surface that overlapped or were in proximity with these projected PdPs were identified as the corresponding PdP. Due to the high morphological variability of this region, the shape of a given PdP varied across participants. However, the relative location of these PdPs appears conserved across individuals given that they are visible on an average cortical surface (**Fig. 7a and 7b**). The PdP posterior to HG was consistently identified across participants and labeled as the PdP-PHG. We quantified the overlap of a participant’s anatomically-defined PdP-PHG with the functionally-defined high-low-high tonotopic map (PHG map) using a simple Boolean operation resulting in a hit if the anatomically-defined PdP-PHG shared any vertices with the functionally-defined PHG map.

**pRF Decoding Analysis:**

From our pRF analysis in the HG ROI, we determined a developmental increase in pRF coverage over low frequencies as well as in pRF tuning width (σ) specifically in low frequency preferring (μ < 150 Hz) voxels. We simulated an adult and pediatric population of pRFs based on experimentally derived pRF properties in this ROI to link pRF development to the decoding of low frequencies. First, we initialized populations of 100 adult-like and 100 child-like pRFs using the group-averaged σ in low μ voxels in HG for adults (278 Hz) and children (210 Hz), respectively. For each population, we generated a grid of biologically plausible μ values and constructed a matrix of 1-D Gaussian pRFs (15 frequency bins × 100 pRFs), with each pRF centered at a different μ value. Least-squares regression was then used to solve for the weights assigned to each pRF that minimized the residuals between the simulated population and the experimentally derived group averaged pRF coverage plot. Preferred frequency (μ) values were subsequently sampled from the initial grid according to the distribution of the fitted weights. The resulting μ values were assigned to each pRF and used to produce a population of 1-D Gaussian pRFs (15 frequency bins x 100 pRFs) that, when averaged across pRFs, best approximated the experimental pRF coverage plots for adults and children. To determine how well a given pRF population could reconstruct a low frequency signal, we estimated the response of each pRF to a 150 Hz tone by evaluating its response at the frequency bin corresponding to 150 Hz. The weighted average of these responses was used to predict the stimulus frequency, and decoding error was determined as the difference between the predicted and actual tone (in Hz). To investigate the effect of noise on pRF decoding, we added increasing levels of Gaussian white noise to the population response, with the standard deviation of the noise set to 1–100% of the mean population response, before calculating the weighted average.

### Statistical Analysis

A linear mixed-effects (LME) model was fit to tone-in-noise detection thresholds from the *Noisy Nest* 3-AFC task, with age, frequency and white noise condition as fixed effects and a random intercept for participant. Statistical analyses comparing pRF parameters between age groups were performed using a two-sample t-test. Each participant contributed a single point representing a given pRF parameter in an ROI (averaged across hemispheres). Two-sample t-tests were similarly used to compare ROI surface area between adults and children. A two-way repeated-measures ANOVA with age and frequency preference as the between- and within-subject factors respectively was utilized to determine developmental differences in σ within low vs high frequency preferring voxels. A two-way repeated-measures ANOVA with age and frequency as the between- and within-subject factors was used to analyze cochleotopic coverage differences between age groups. We used a two-way repeated-measures ANOVA with ROI and frequency as within-subject factors to assess differences in cochleotopic coverage between two ROIs within each age group. A LME model was fit to tone-in-noise detection thresholds to determine the behavioral relevance of our neural results: age, frequency and cochleotopic coverage were treated as fixed effects and individual slopes were treated as random effects. Another LME model was used to assess the effect of age group and noise on the frequency decoding error of simulated pRF populations, with age and noise factor as the fixed effects and simulation iteration as the random effect. Structure-function coupling analysis was performed using a Boolean operation score of 1 for a hit of any shared vertices between anatomically- and functionally-defined ROIs and 0 for no overlapping vertices. The degrees of freedom in all analyses are reported.

## Data and Code Availability

Data visualization was performed using custom-made scripts in MATLAB (https://github.com/damolaogunlade/tonotopy_figures)

## Author Contribution

OO contributed to study design, data collection, analysis, and manuscript preparation. JG contributed to study design, analysis, and manuscript preparation.

## Acknowledgments

This material is based upon work supported by the National Science Foundation under CAREER Award No. 2337373, by the Whitehall Foundation (Award no. 2023-08-052), and funds from Princeton University to JG. Funding for the acquisition of the data was provided in part by the Regina and John Scully ’66 Center for the Neuroscience of Mind and Behavior. NIH grant T32MH065214 was used to support Oyindamola Ogunlade while they were a graduate student at Princeton University.

---

**Supplementary Figure 1:**
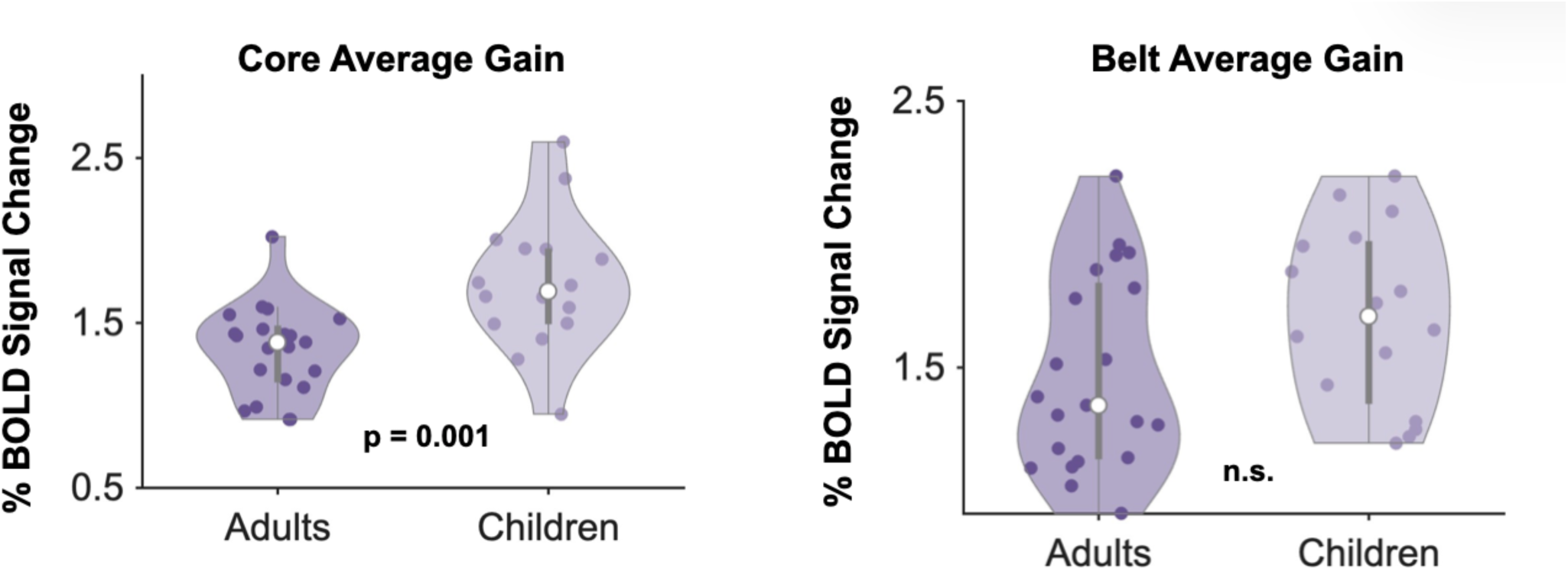
The pRF gain parameter in the classically-defined core and posterolateral belt. Children (light purple) showed larger average gain across pRFs within the core (t(35) = −3.62, p = 0.001) compared to adults (dark purple). There was no significant difference from adults within the posterolateral belt (t(35) = −1.88, p = 0.07; Fig S1), and gain did not vary as a function of frequency and age (ANOVA interaction: F(l, 141) = 1.17, p = 0.28).

**Supplementary Figure 2:**
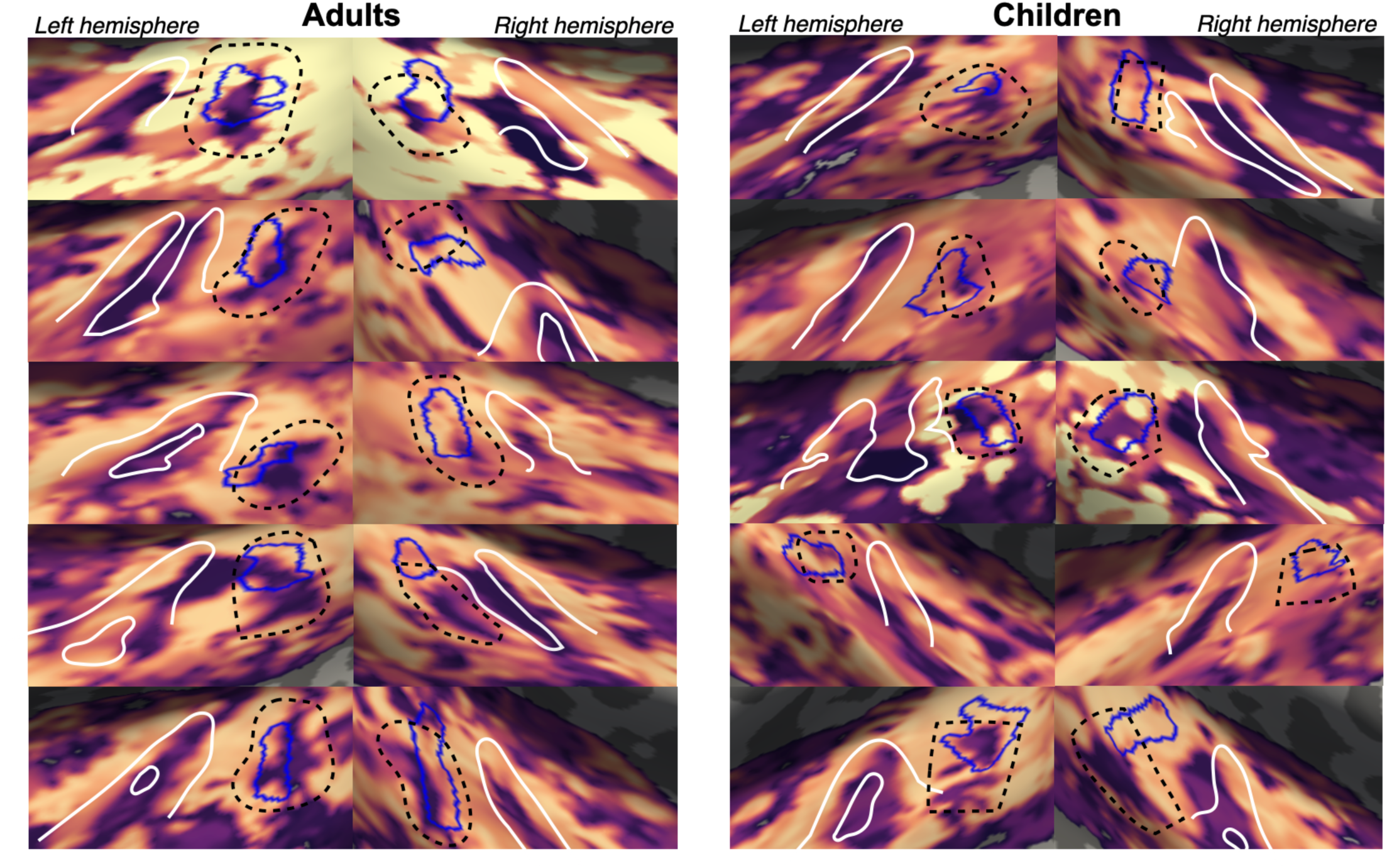
**The high-low-high frequency gradient posterior to HG overlapping with a *plfy de passage.*** The tonotopic frequency gradient is outlined in dotted black in adults (left) and children (right) in both hemispheres. Each row is a participant, showing their left and right superior temporal cortex on the left and right side, respectively. The *p/i de passage* is outlined in solid blue. Hes.chl’.s Gyrus is outlined in solid white.

## References

1. Formisano, E. et al. Mirror-Symmetric Tonotopic Maps in Human Primary Auditory Cortex. Neuron 40, 859–869 (2003).

2. Talavage, T. M. et al. Tonotopic Organization in Human Auditory Cortex Revealed by Progressions of Frequency Sensitivity. J. Neurophysiol. 91, 1282–1296 (2004).

3. Humphries, C., Liebenthal, E. & Binder, J. R. Tonotopic organization of human auditory cortex. NeuroImage 50, 1202–1211 (2010).

4. Costa, S. D. et al. Human Primary Auditory Cortex Follows the Shape of Heschl’s Gyrus. J. Neurosci. 31, 14067–14075 (2011).

5. Baumann, S., Petkov, C. I. & Griffiths, T. D. A unified framework for the organization of the primate auditory cortex. Front. Syst. Neurosci. 7, (2013).

6. Leaver, A. M. & Rauschecker, J. P. Cortical Representation of Natural Complex Sounds: Effects of Acoustic Features and Auditory Object Category. J. Neurosci. 30, 7604–7612 (2010).

7. Staeren, N., Renvall, H., Martino, F. D., Goebel, R. & Formisano, E. Sound Categories Are Represented as Distributed Patterns in the Human Auditory Cortex. Curr. Biol. 19, 498–502 (2009).

8. Hamilton, L. S., Oganian, Y., Hall, J. & Chang, E. F. Parallel and distributed encoding of speech across human auditory cortex. Cell 184, 4626–4639.e13 (2021).

9. Davis, M. H. & Johnsrude, I. S. Hierarchical Processing in Spoken Language Comprehension. J. Neurosci. 23, 3423–3431 (2003).

10. Wessinger, C. M. et al. Hierarchical organization of the human auditory cortex revealed by functional magnetic resonance imaging. J. Cogn. Neurosci. 13, 1–7 (2001).

11. Heer, W. A. de, Huth, A. G., Griffiths, T. L., Gallant, J. L. & Theunissen, F. E. The Hierarchical Cortical Organization of Human Speech Processing. J. Neurosci. 37, 6539–6557 (2017).

12. Functional Organization of the Visual Cortex. in Progress in Brain Research vol. 58 209–218 (Elsevier, 1983).

13. Engel, S. Retinotopic organization in human visual cortex and the spatial precision of functional MRI. Cereb. Cortex 7, 181–192 (1997).

14. Tootell, R. B. H. et al. Functional analysis of primary visual cortex (V1) in humans. Proc. Natl. Acad. Sci. 95, 811–817 (1998).

15. Arcaro, M. J., McMains, S. A., Singer, B. D. & Kastner, S. Retinotopic Organization of Human Ventral Visual Cortex. J. Neurosci. 29, 10638–10652 (2009).

16. Witthoft, N. et al. Where is human V4? Predicting the location of hV4 and VO1 from cortical folding. Cereb. Cortex N. Y. N 1991 24, 2401–2408 (2014).

17. Benson, N. C. & Winawer, J. Bayesian analysis of retinotopic maps. eLife 7, e40224 (2018).

18. Gomez, J. et al. Development of population receptive fields in the lateral visual stream improves spatial coding amid stable structural-functional coupling. NeuroImage 188, 59–69 (2019).

19. Gomez, J., Natu, V., Jeska, B., Barnett, M. & Grill-Spector, K. Development differentially sculpts receptive fields across early and high-level human visual cortex. Nat. Commun. 9, 788 (2018).

20. Merzenich, M. M. & Brugge, J. F. Representation of the cochlear partition on the superior temporal plane of the macaque monkey. Brain Res. 50, 275–296 (1973).

21. Merzenich, M. M., Knight, P. L. & Roth, G. L. Representation of cochlea within primary auditory cortex in the cat. J. Neurophysiol. 38, 231–249 (1975).

22. Reale, R. A. & Imig, T. J. Tonotopic organization in auditory cortex of the cat. J. Comp. Neurol. 192, 265–291 (1980).

23. Morel, A., Garraghty, P. E. & Kaas, J. H. Tonotopic organization, architectonic fields, and connections of auditory cortex in macaque monkeys. J. Comp. Neurol. 335, 437–459 (1993).

24. Brodmann, K. Vergleichende Lokalisationslehre Der Grosshirnrinde in Ihren Prinzipien Dargestellt Auf Grund Des Zellenbaues. (1909).

25. Economo, C. von, Koskinas, G. N. & Triárchou, L. K. Atlas of Cytoarchitectonics of the Adult Human Cerebral Cortex: Compiled at the Psychiatric Clinic of Hofrat Julius Wagner von Jauregg, Vienna. (Karger, Basel, 2008).

26. EconomoC. F. von & Koskinas, G. N. Die Cytoarchitektonik der Hirnrinde des erwachsenen Menschen. (J. Springer, 1925).

27. Hackett, T. A. Anatomic organization of the auditory cortex. in Handbook of Clinical Neurology vol. 129 27–53 (Elsevier, 2015).

28. Lauter, J. L., Herscovitch, P., Formby, C. & Raichle, M. E. Tonotopic organization in human auditory cortex revealed by positron emission tomography. Hear. Res. 20, 199–205 (1985).

29. Moerel, M., De Martino, F. & Formisano, E. An anatomical and functional topography of human auditory cortical areas. Front. Neurosci. 8, (2014).

30. Rauschecker, J. P., Tian, B. & Hauser, M. Processing of Complex Sounds in the Macaque Nonprimary Auditory Cortex. Science 268, 111–114 (1995).

31. Saenz, M. & Langers, D. R. M. Tonotopic mapping of human auditory cortex. Hear. Res. 307, 42–52 (2014).

32. Langers, D. R. M. Assessment of tonotopically organised subdivisions in human auditory cortex using volumetric and surface-based cortical alignments. Hum. Brain Mapp. 35, 1544–1561 (2013).

33. Müller, R., Fleischer, G. & Schneider, J. Pure-tone auditory threshold in school children. Eur. Arch. Otorhinolaryngol. 269, 93–100 (2012).

34. Trehub, S. E., Schneider, B. A., Morrongiello, B. A. & Thorpe, L. A. Auditory sensitivity in school-age children. J. Exp. Child Psychol. 46, 273–285 (1988).

35. Allen, P., Wightman, F., Kistler, D. & Dolan, T. Frequency Resolution in Children. J. Speech Lang. Hear. Res. 32, 317–322 (1989).

36. Irwin, R. J., Stillman, J. A. & Schade, A. The width of the auditory filter in children. J. Exp. Child Psychol. 41, 429–442 (1986).

37. Rubensohn. Mastoid Pneumatization in Children at Various Ages. Acta Otolaryngol. (Stockh.) 10.3109/00016486509126983 (1965) doi:10.3109/00016486509126983.

38. Moore, J. K. & Linthicum Jr., F. H. The human auditory system: A timeline of development. Int. J. Audiol. 46, 460–478 (2007).

39. Wunderlich, J. L., Cone-Wesson, B. K. & Shepherd, R. Maturation of the cortical auditory evoked potential in infants and young children. Hear. Res. 212, 185–202 (2006).

40. Jones, M. K. et al. Auditory Processing Differences in Toddlers With Autism Spectrum Disorder. J. Speech Lang. Hear. Res. 63, 1608–1617 (2020).

41. Alcántara, J. I., Weisblatt, E. J. L., Moore, B. C. J. & Bolton, P. F. Speech-in-noise perception in high-functioning individuals with autism or Asperger’s syndrome. J. Child Psychol. Psychiatry 10.1111/j.1469-7610.2004.t01-1-00303.x (2004) doi:10.1111/j.1469-7610.2004.t01-1-00303.x.

42. Lepistö, T. et al. Auditory stream segregation in children with Asperger syndrome. Biol. Psychol. 82, 301–307 (2009).

43. Russo, N., Nicol, T., Trommer, B., Zecker, S. & Kraus, N. Brainstem transcription of speech is disrupted in children with autism spectrum disorders. Dev. Sci. 12, 557–567 (2009).

44. Koravand, A., Jutras, B. & Lassonde, M. Abnormalities in cortical auditory responses in children with central auditory processing disorder. Neuroscience 346, 135–148 (2017).

45. Purdy, Kelly & Davies. Auditory Brainstem Response, Middle Latency Response, and Late Cortical Evoked Potentials in Children with Learning Disabilities. J. Am. Acad. Audiol. 10.1055/s-0040-1715981 (2002) doi:10.1055/s-0040-1715981.

46. Rance & Tomlin. Maturation of the Central Auditory Nervous System in Children with Auditory Processing Disorder. Semin. Hear. 10.1055/s-0035-1570328 (2016) doi:10.1055/s-0035-1570328.

47. Himmelberg, M. M. et al. Comparing retinotopic maps of children and adults reveals a late-stage change in how V1 samples the visual field. Nat. Commun. 14, 1561 (2023).

48. Sydnor, V. J. et al. Neurodevelopment of the association cortices: Patterns, mechanisms, and implications for psychopathology. Neuron 109, 2820–2846 (2021).

49. Burt, J. B. et al. Hierarchy of transcriptomic specialization across human cortex captured by structural neuroimaging topography. Nat. Neurosci. 21, 1251–1259 (2018).

50. Gomez, J., Zhen, Z. & Weiner, K. S. Human visual cortex is organized along two genetically opposed hierarchical gradients with unique developmental and evolutionary origins. PLoS Biol. 17, e3000362 (2019).

51. Paquola, C. et al. Shifts in myeloarchitecture characterise adolescent development of cortical gradients. eLife 8, e50482.

52. Hill, J. et al. Similar patterns of cortical expansion during human development and evolution. Proc. Natl. Acad. Sci. 107, 13135–13140 (2010).

53. Sowell, E. R. et al. Longitudinal Mapping of Cortical Thickness and Brain Growth in Normal Children. J. Neurosci. 24, 8223–8231 (2004).

54. Da Costa, S., Van Der Zwaag, W., Miller, L. M., Clarke, S. & Saenz, M. Tuning In to Sound: Frequency-Selective Attentional Filter in Human Primary Auditory Cortex. J. Neurosci. 33, 1858–1863 (2013).

55. De Martino, F. et al. High-Resolution Mapping of Myeloarchitecture In Vivo: Localization of Auditory Areas in the Human Brain. Cereb. Cortex N. Y. NY 25, 3394–3405 (2015).

56. Morosan, P. et al. Human Primary Auditory Cortex: Cytoarchitectonic Subdivisions and Mapping into a Spatial Reference System. NeuroImage 13, 684–701 (2001).

57. Thomas, J. M. et al. Population receptive field estimates of human auditory cortex. NeuroImage 105, 428–439 (2015).

58. Dumoulin, S. O. & Wandell, B. A. Population receptive field estimates in human visual cortex. NeuroImage 39, 647–660 (2008).

59. Mann, V. A., Diamond, R. & Carey, S. Development of voice recognition: Parallels with face recognition. J. Exp. Child Psychol. 27, 153–165 (1979).

60. Watson, A. B. QUEST+: A general multidimensional Bayesian adaptive psychometric method. J. Vis. 17, 10 (2017).

61. Fletcher, H. Auditory Patterns. Rev. Mod. Phys. 12, 47–65 (1940).

62. Power, J. D., Barnes, K. A., Snyder, A. Z., Schlaggar, B. L. & Petersen, S. E. Spurious but systematic correlations in functional connectivity MRI networks arise from subject motion. NeuroImage 59, 2142–2154 (2012).

63. Leonard, C. M., Puranik, C., Kuldau, J. M. & Lombardino, L. J. Normal variation in the frequency and location of human auditory cortex landmarks. Heschl’s gyrus: where is it? Cereb. Cortex 8, 397–406 (1998).

64. Glasser, M. F. & Essen, D. C. V. Mapping Human Cortical Areas In Vivo Based on Myelin Content as Revealed by T1- and T2-Weighted MRI. J. Neurosci. 31, 11597–11616 (2011).

65. Preziosa, P. et al. Neurite density explains cortical T1-weighted/T2-weighted ratio in multiple sclerosis. J. Neurol. Neurosurg. Psychiatry 92, 790–792 (2021).

66. Grill-Spector, K. & Weiner, K. S. The functional architecture of the ventral temporal cortex and its role in categorization. Nat. Rev. Neurosci. 15, 536–548 (2014).

67. Rosenke, M. et al. A cross-validated cytoarchitectonic atlas of the human ventral visual stream. NeuroImage 170, 257–270 (2018).

68. Gratiolet, P.-L. (1815-1865) A. du texte. Mémoire Sur Les Plis Cérébraux de l’homme et Des Primatès [”sic”]. ATLAS / Par M. Pierre Gratiolet,… (1834).

69. Mangin, J.-F. et al. “Plis de passage” Deserve a Role in Models of the Cortical Folding Process. Brain Topogr. 32, 1035–1048 (2019).

70. Cachia, A. et al. How interindividual differences in brain anatomy shape reading accuracy. Brain Struct. Funct. 223, 701–712 (2018).

71. Kay, K. N., Weiner, K. S. & Grill-Spector, K. Attention Reduces Spatial Uncertainty in Human Ventral Temporal Cortex. Curr. Biol. 25, 595–600 (2015).

72. Okada, K., Venezia, J. H., Matchin, W., Saberi, K. & Hickok, G. An fMRI Study of Audiovisual Speech Perception Reveals Multisensory Interactions in Auditory Cortex. PLOS ONE 8, e68959 (2013).

73. Calvert, G. A. & Campbell, R. Reading speech from still and moving faces: The neural substrates of visible speech. J. Cogn. Neurosci. 15, 57–70 (2003).

74. Balk, M. et al. Synchrony of audio-visual speech stimuli modulates left superior temporal sulcus. Neuroreport 21, 822–6 (2010).

75. Martuzzi, R. et al. Multisensory Interactions within Human Primary Cortices Revealed by BOLD Dynamics. Cereb. Cortex 17, 1672–1679 (2007).

76. Molholm, S. et al. Multisensory auditory-visual interactions during early sensory processing in humans: a high-density electrical mapping study. Brain Res. Cogn. Brain Res. 14, 115–128 (2002).

77. Battal, C., Rezk, M., Mattioni, S., Vadlamudi, J. & Collignon, O. Representation of Auditory Motion Directions and Sound Source Locations in the Human Planum Temporale. J. Neurosci. 39, 2208–2220 (2019).

78. Hackett, T. A., Preuss, T. M. & Kaas, J. H. Architectonic identification of the core region in auditory cortex of macaques, chimpanzees, and humans. J. Comp. Neurol. 441, 197–222 (2001).

79. Lage-Castellanos, A., De Martino, F., Ghose, G. M., Gulban, O. F. & Moerel, M. Selective attention sharpens population receptive fields in human auditory cortex. Cereb. Cortex N. Y. NY 33, 5395–5408 (2022).

80. Glasberg, B. R. & Moore, B. C. Derivation of auditory filter shapes from notched-noise data. Hear. Res. 47, 103–138 (1990).

81. Bodin, C. et al. Plis de passage in the superior temporal sulcus: Morphology and local connectivity. NeuroImage 225, 117513 (2021).

82. Wandell, B. A. & Winawer, J. Computational neuroimaging and population receptive fields. Trends Cogn. Sci. 19, 349–357 (2015).

83. Bonte, M. et al. Development from childhood to adulthood increases morphological and functional inter-individual variability in the right superior temporal cortex. NeuroImage 83, 739–750 (2013).

84. Berg, Fuchs, Wirkner, Loeffler & Et., A. The Speaking Voice in the General Population: Normative Data and Associations to Sociodemographic and Lifestyle Factors. J. Voice 10.1016/j.jvoice.2016.06.001 (2017) doi:10.1016/j.jvoice.2016.06.001.

85. Moerel, M., Martino, F. D. & Formisano, E. Processing of Natural Sounds in Human Auditory Cortex: Tonotopy, Spectral Tuning, and Relation to Voice Sensitivity. J. Neurosci. 32, 14205–14216 (2012).

86. Langner, G., Sams, M., Heil, P. & Schulze, H. Frequency and periodicity are represented in orthogonal maps in the human auditory cortex: evidence from magnetoencephalography. J. Comp. Physiol. A 181, 665–676 (1997).

87. Peelle, J. E. Methodological challenges and solutions in auditory functional magnetic resonance imaging. Front. Neurosci. 8, 253 (2014).

88. ISO-226-2003.pdf.

89. Fischl, B. FreeSurfer. NeuroImage 62, 774–781 (2012).

90. Fischl, B., Sereno, M. I., Tootell, R. B. H. & Dale, A. M. High-resolution intersubject averaging and a coordinate system for the cortical surface. Hum. Brain Mapp. 8, 272–284 (1999).

91. Dale, A. M., Fischl, B. & Sereno, M. I. Cortical surface-based analysis. I. Segmentation and surface reconstruction. NeuroImage 9, 179–194 (1999).

